# The increased hinge flexibility of an IgG1-IgG3 hybrid monoclonal enhances Fc-mediated protection against group A streptococci

**DOI:** 10.1101/2023.10.14.562368

**Authors:** Arman Izadi, Yasaman Karami, Eleni Bratanis, Sebastian Wrighton, Hamed Khakzad, Maria Nyblom, Berit Olofsson, Lotta Happonen, Di Tang, Michael Nilges, Johan Malmström, Wael Bahnan, Oonagh Shannon, Lars Malmström, Pontus Nordenfelt

## Abstract

Antibodies are central to the immune response against microbes. We have previously generated a protective IgG1 monoclonal antibody targeting the M protein, a critical virulence factor of *Streptococcus pyogenes.* Here, we generated this antibody in all human IgG subclasses and evaluated their function. Despite significantly reduced binding, the IgG3 subclass antibody demonstrated remarkably enhanced opsonic function. We hypothesized that increased Fc flexibility could explain this improved efficacy. We engineered a hybrid IgG subclass antibody, IgGh, containing the backbone of IgG1 with the hinge of IgG3, leaving the Fabs unchanged. The IgGh maintained a similar binding ability as IgG1 while gaining the strong opsonic function seen with IgG3. Molecular dynamics simulations of the different antibodies showed altered IgG Fab-antigen interactions, reflecting the differences observed in affinity. More importantly, when the antibodies were bound to the antigen, the simulations showed that the Fc of both IgGh and IgG3 exhibited extensive movement in 3D space relative to the M protein. The increased flexibility of IgGh directly translated to enhanced opsonic function and significantly increased the protection against infection with *Streptococcus pyogenes* in mice. Our findings demonstrate how altering Fc flexibility can improve Fc-mediated opsonic function and how modifications in the constant domain can regulate Fab-antigen interactions. In addition, the enhanced *in vivo* function of a more flexible IgG provides new therapeutic opportunities for monoclonal antibodies.

**One sentence summary:** Antibody Fc flexibility in 3D space correlates with efficient Fc-mediated phagocytosis of streptococci

## Introduction

The immune system generates antibodies against invading pathogens as an essential part of the adaptive immune response. VDJ gene recombination in B cells defines the basic antigen-binding properties of antibodies (1,2), and B cells selected within the lymphoid tissues undergo class-switching and somatic hypermutation. The classical model of antibody variable and constant domain independence describes that class-switching only alters antibody effector function and does not alter affinity (3). However, some findings suggest that the constant domain could influence the binding to antigens (4–7), providing a potential mechanism where subclass and class-switching add further affinity diversity in addition to VDJ-recombination and somatic hypermutation (4–5).

The human IgG antibody class contains four subclasses with varying abilities to trigger effector functions such as complement activation, phagocytosis, and cell cytotoxicity. These immune activities are mediated through binding to Fc-receptors on the surface of immune cells or binding to soluble complement proteins through the Fc-portion (1). These subclasses differ in the constant region of the heavy chain. Major differences between the subclasses lie in the hinge region that connects the Fab to the Fc domains (Fig. 1A). Although there are at least 13 allotypes of the IgG3 subclass (1), a common trait is that they have the most extended hinge region. This increased hinge length makes them more flexible, while the shorter hinge region of IgG2 is the most rigid (8). Furthermore, IgG3 is most efficient at activating the classical complement pathway (9). IgG3 and IgG1 have similar affinities to Fc-gamma receptors in humans, while IgG2 and IgG4 overall have a lower receptor affinity (10).

**Figure 1.**
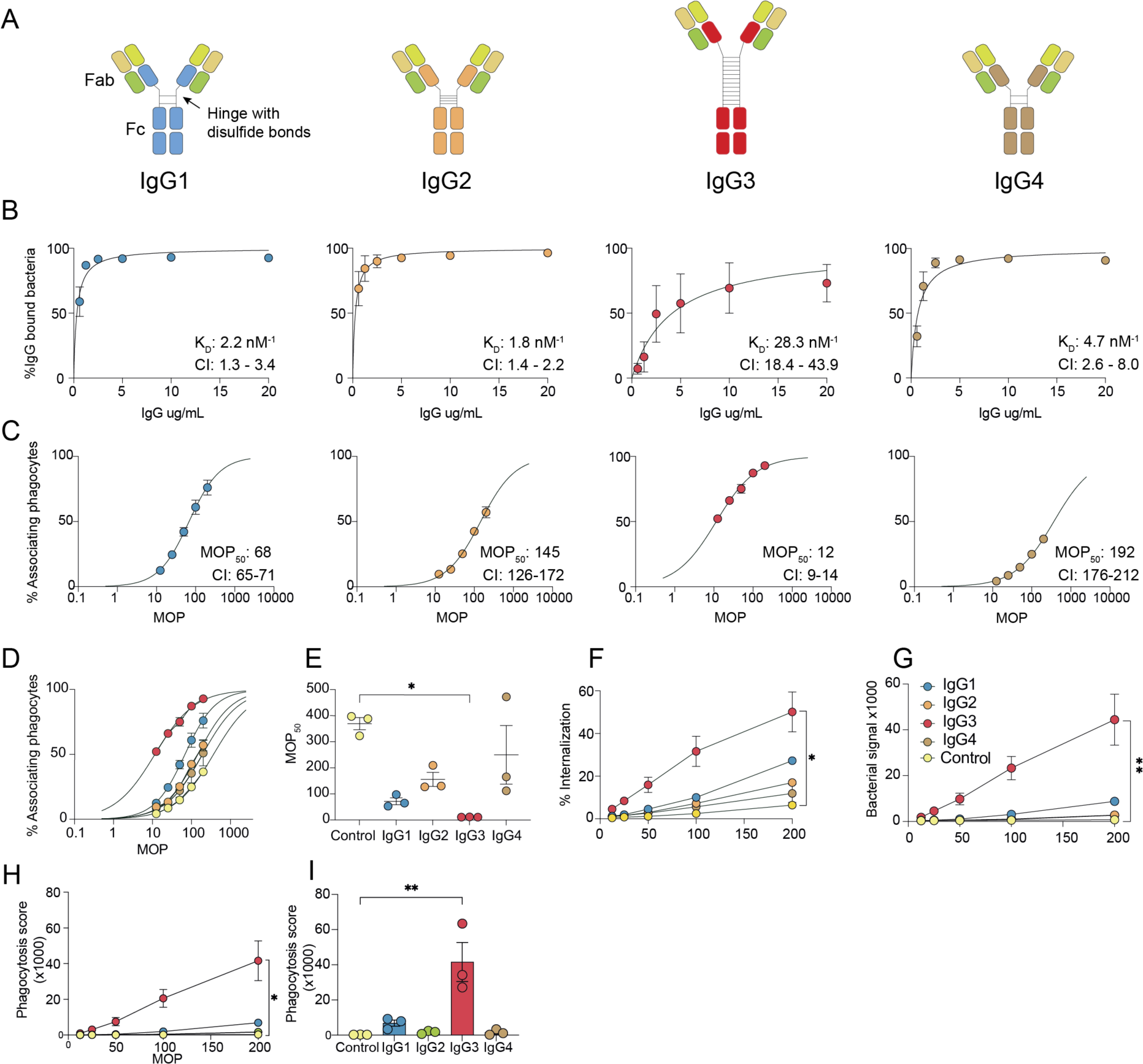
IgG3 exhibits potent phagocytosis efficacy despite reduced binding to antigen. **A** The four subclasses of Ab25. The antibodies consist of two identical heavy chains and identical light chains. The variable heavy domain (olive green), the constant domain of the light chain (green), and the variable light domain (beige) are identical for the four antibodies.The constant domains of the heavy chains are depicted in different colors: blue for IgG1, orange for IgG2, red for IgG3, and brown for IgG4. **B** Affinity curves for the IgG subclasses. The Y-axis is the percentage of GFP-expressing bacteria positive for antibodies. The K_D_ values (nM^-1^) with a 95% confidence interval are shown in each respective graph. Data points are from three independent experiments. **C** MOP_50_ curves of the subclasses, bacteria were opsonized with 15 ug/mL of each respective antibody or control. The Y-axis shows the percentage of THP-1 cells that are positive for bacteria, while the X-axis depicts the ratio of bacteria to phagocytes added (MOP). The MOP_50_ values with a 95% confidence interval are shown in each graph. The data points are from three independent experiments. **D** Shows the MOP_50_ curves of all the antibody treatments in one graph. Control is Xolair, human IgG1 anti-IgE. **E** Individual MOP_50_ values from the three independent experiments. **F** Percentage of THP-1 cells with internalized bacteria across the MOP range. **G** Amount of bacteria that are phagocytosed by the whole THP-1 cell population measured as median MFI of FITC-A (bacterial signal, Oregon Green). **H-I** Phagocytosis score for each antibody treatment is shown with MOP on the X-axis for **H**, while in **I**, the MOP is 200. All statistical analysis was done, comparing the treatments to the control antibody with Kruskal-Wallis multiple comparisons corrected by Dunn’s correction test. * denotes P-value < 0.05 and P-value > 0.05 is ns. The data points in figure **B-I** represent the mean value, and the error bars are in SEM.

We have previously generated one IgG1 mAb (Ab25) against the M protein of *Streptococcus pyogenes* (11). *S. pyogenes* has several virulence factors, of which the M protein is one of the most studied due to its importance for virulence (12). It is a cell-wall-attached protein with a dimeric coiled-coil structure that extends out from the surface of the bacteria (12). The M protein has several immune evasive mechanisms, such as binding antibodies via their Fc-portion, which can inhibit Fc-mediated phagocytosis (13–14) or reducing phagocytosis through antibody-induced fibronectin binding (15). In addition, the M protein inhibits complement-mediated phagocytosis by sequestering C4BP and fibrinogen binding (16). These mechanisms are important for *S. pyogenes* virulence and survival in the host. M protein is a target for vaccine development against *S. pyogenes*. However, no vaccines have been approved for clinical use due to a lack of efficacy or because they promote the emergence of autoreactive antibodies (17). The generation of distinct mAbs against *S. pyogenes* could be a viable therapeutic alternative. Indeed, the IgG1 Ab25 mAb was protective when used as a prophylactic in a mouse subcutaneous infection (11). We have previously observed that enriched binding of polyclonal IgG3 antibodies to M protein was linked to enhanced phagocytosis (18). However, the importance of subclass for effective function against *S. pyogenes* or other bacterial pathogens is not well understood.

In this work, we generated all four human subclasses of Ab25 and characterized them *in vitro*. We observed that the IgG3 subclass of Ab25 had reduced binding to M1 protein as compared to the other subclasses. Still, this antibody subclass had several-fold higher opsonic efficacy than the original IgG1 Ab25. We hypothesized that the IgG3 subclass efficacy could be explained by its more flexible hinge region. To analyze hinge flexibility, we exchanged the original IgG1 to contain the IgG3 hinge instead. Molecular dynamics (MD) simulations showed that this new hybrid mAb (IgGh) and the IgG3 mAb display much higher flexibility than the original IgG1. The constant domain also changed how the Fabs interacted with the antigen. The IgGh showed a similar binding strength as IgG1 but exceeded both IgG1 and IgG3 in opsonic efficacy. Our *in vitro* findings translated to an *in vivo* infection model in mice where the IgGh exceeded its parent subclasses with regard to protection during systemic infection. We, therefore, propose that increasing antibody flexibility through hinge engineering has promising therapeutic implications.

## Results

### IgG3 exhibits potent phagocytosis efficacy despite reduced binding to antigen

Previously, we generated a human IgG1 antibody, Ab25, against the M protein of *Streptococcus pyogenes* which was opsonic *in vitro* and protective *in vivo* (11). This antibody was generated using a human B cell from a convalescent donor. The original subclass is not known. Using PCR and homologous recombination, we exchanged the subclass-specific domains of the heavy chain, generating three new plasmids encoding the remaining subclasses IgG2, IgG3, and IgG4 (**Fig. 1A**). The hinge length varies, with IgG1 having a 15 aa flexible hinge, IgG2 having a 12 aa rigid hinge (due to more disulfide bridges), and IgG4 having a 12 aa flexible hinge. IgG3 has about four times longer hinge region, distributed across 13 different allotypes with up to 62 aa hinge length (1), and for this study, we used allotype IGHG3*11 with a 62 aa hinge.

In previous work, we attempted to measure the affinity of Ab25 IgG1 to both purified and recombinant M protein by using surface plasmon resonance (SPR). These attempts were unsuccessful. This is most likely due to the M protein’s conformational stability being highly dependent on temperature, dimerization, and being attached to its native bacterial surface (19). Instead, we measured the direct functional affinity (likely equivalent to avidity) of antibodies to M protein on live bacteria (**Supp. Fig. 1A**) (11). We calculated the dissociation constant (K_D_) for all Ab25 subclasses using a non-linear regression analysis. This analysis revealed different affinity profiles across the subclasses (**Fig. 1B**). IgG1, IgG2, and IgG4 showed similar affinities (IgG1 K_D_ = 2.2; IgG2 KD = 1.8; IgG4 KD 4.7 nM^-1,^ respectively). IgG3 showed more than a 14-fold decrease in affinity compared to IgG1 (IgG3 K_D_ = 28.3 nM^-1^). Our results indicate that subclass-switching Ab25 IgG1 to IgG3 leads to reduced binding to the M protein, while for IgG2 and IgG4, binding affinity is retained.

Having established the binding properties of the Ab25 IgG subclass variants, we investigated their functional output in terms of Fc-mediated phagocytosis. We used a previously established opsonophagocytosis assay for *S. pyogenes* with heat-killed SF370 bacteria and human phagocytes of the monocytic THP-1 cell line (11, 20). The opsonic ability of the subclasses was assessed by varying the ratio of prey to phagocyte (multiplicity of prey, MOP). We calculated the number of bacteria relative to phagocytes needed to elicit a 50% association (MOP_50_) (20). A lower MOP_50_ value means it is more efficient at mediating bacteria-monocyte interaction.

The MOP_50_ analysis showed that IgG3 exhibited a potent opsonic phenotype (**Fig. 1C**). There was a 6-fold increase in opsonic ability (MOP_50_ = 12) compared to the original Ab25 IgG1 (MOP_50_ = 68) and a 12-16-fold increase compared to IgG2 (MOP_50_ = 145) and IgG4 (MOP_50_ = 192) (**Fig. 1D-E**). To put these values into context, the negative Fc-control exhibited a 30-fold lower efficacy compared to IgG3 (MOP_50_ = 367). We analyzed the percentage of phagocytes with internalized bacteria and saw a similar outcome as the MOP_50_ analysis with the following order IgG3>IgG1>IgG2>IgG4. (**Fig. 1F**). Additionally, when considering the bacterial signal, a quantity metric, we assessed the relative quantity of bacteria phagocytosed (**Supp. Fig. 1B-C**). This analysis shows that IgG3 promotes the most efficient phagocyte-bacteria interaction (**Fig. 1G**). Finally, we calculated a phagocytosis score based on the combined phagocyte association and bacterial signal (MFI) of the THP-1 population. This analysis shows that IgG3 is more potent than the other subclasses, with a 7-fold higher score than IgG1 at MOP 200 (41600 vs. 6900, **Fig.1H-I**), 26-fold higher than both IgG2 and IgG4 (1600 respectively) and 130-fold higher than control (300). The four assessed metrics (MOP_50_, internalization, bacterial signal, and phagocytosis score) showed that Ab25 IgG3, despite its reduced reactivity to M1, is a more potent opsonizing antibody than the other subclasses.

### The IgG3 hinge provides broad Fc spatial mobility spectra and flexibility relative to M protein

Our *in vitro* findings of increased opsonic ability of IgG3 align with other recent findings with increased Fc-effector functions of this subclass (21–23). However, we observed a discrepancy with lower binding yet stronger functional output. We hypothesized that this increased functionality is due to the extended hinge region of IgG3. To investigate this, we engineered a hybrid mAb containing the backbone of IgG1 but with the hinge of IgG3 (IgGh). Affinity measurements with directly labeled IgG1 and IgGh antibodies showed that the antigen binding appeared to be unaffected by this modification **(Supp. Fig. 2B)**. To assess if the hinge of IgG3 grants increased Fc flexibility, we performed three replicates of 1 µs molecular dynamics (MD) simulations for IgG1, IgG3, and the IgGh (see Methods section for the details of simulations and **Supplemental Movies 1-9)**. We determined a crystal structure of the Ab25 Fab to enhance the accuracy of the simulations (**Supp. Table 1**, PDB ID: 8C67). We have previously found that the Ab25 binds with both Fabs to two different epitopes on the M protein, so-called dual-Fab cis binding (11). The data obtained by protein cross-linking mass spectrometry was supported by experimental validation showing that the single Fabs of Ab25 IgG1 could barely bind to M protein compared to F(ab’)_2_ fragments (11). Our current MD analysis of the Fab-M1 interactions thus focused on this dual-Fab cis interaction.

To evaluate the stability of the generated trajectories, we measured the root mean square deviations (RMSD) of different domains (M1 protein, Fc, Fab1, and Fab2) with respect to the equilibrated starting structure (**Fig. 2 A-D**). The three systems of IgG1, IgG3, and IgGh represent similar patterns of RMSD within the Fc domain (average values of 5.34±0.88, 4.35±0.91 and 5.28±0.84 Å, respectively), and Fab2 region (average RMSD of 4.89±0.89, 5.16±0.77 and 4.43±0.8 Å, respectively, **Supp. Table 2**). The Fab1 domain of the IgG1 system showed lower RMSD values (average of 3.72±0.64) with respect to the Fab1 of both the IgG3 and IgGh (average values of 5.76±1.06 and 6.49±1.48 Å, respectively). We also measured the root mean square fluctuations (RMSF) of every residue within different domains with respect to the average conformation (**Fig. 2 E-H**). Both IgG3 and IgGh exhibited larger fluctuations of Fab1 (average RMSF of 2.18±1.16 and 2.32±1.33 Å, respectively) compared to IgG1 (average RMSF of 1.92±0.86 Å), hinting at weaker binding to M1 for this Fab (**Supp. Table 2)**. More interestingly, IgGh displayed larger fluctuations and flexibility in its Fc region (average RMSF of 2.88±1.68 Å) compared to both IgG1 and IgG3 (average RMSF of 1.64±1.01 and 1.63±0.96 Å, respectively). Surprisingly IgGh showed smaller fluctuations within Fab2 (average RMSF of 1.62±0.65 Å) compared to both IgG1 and IgG3 (average RMSF of 2.55±0.94 and 2.57±1.24 Å, respectively).

**Figure 2.**
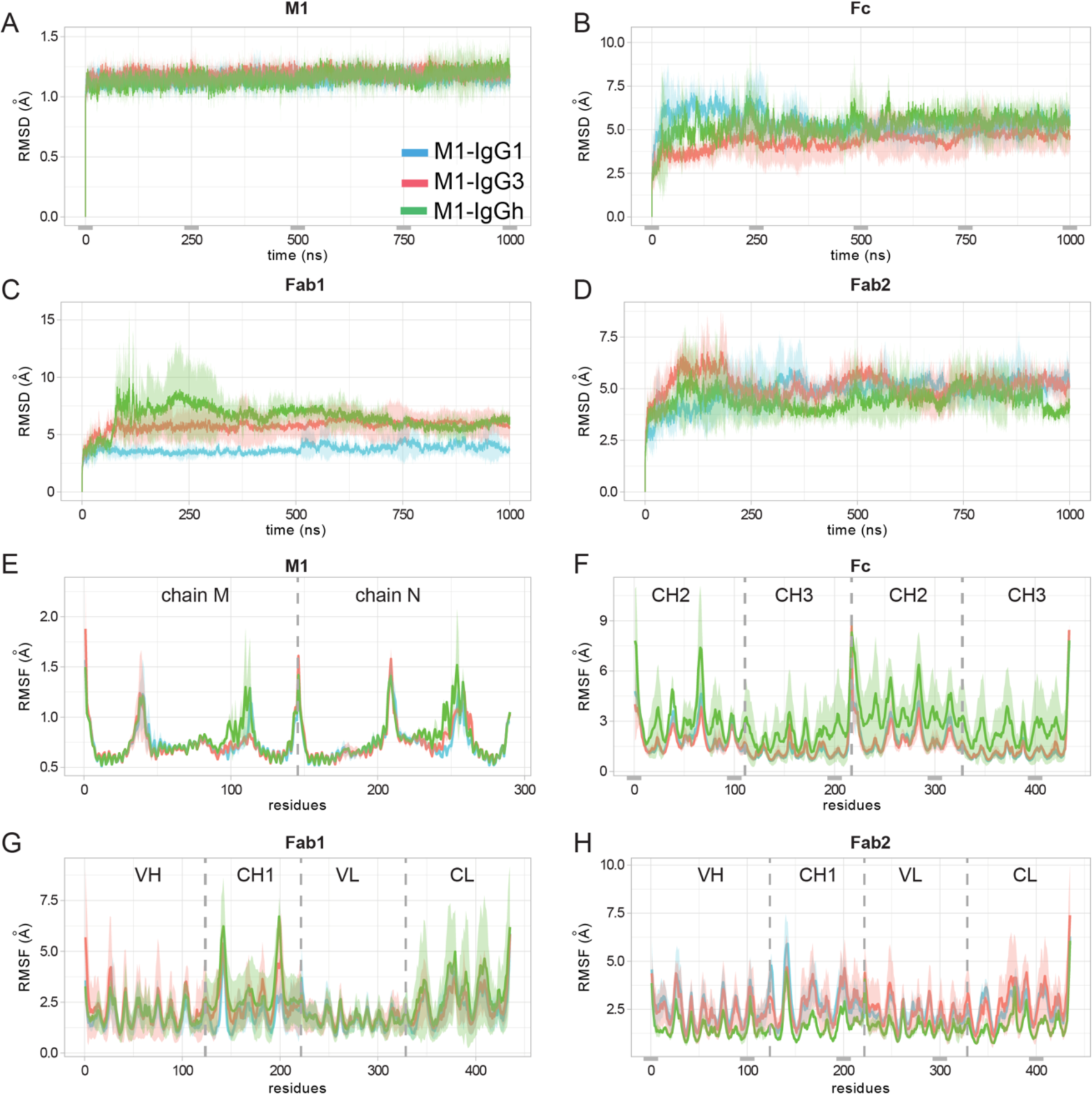
The root mean square deviations and fluctuations for M1-IgGs. The RMSD from the equilibrated structure was computed on the C& atoms of each replicate and every domain: **a)** M1, **b)** Fc, **c)** Fab1, and **d)** Fab2. The RMSF was measured on the C& atoms with respect to the average conformation and averaged by residue, considering the last 900 ns of the MD simulations for the **e)** M1, **f)** Fc, **g)** Fab1, and **h)** Fab2 domains. The average values over the three replicates of each domain are reported in pink, cyan, and green for the M1-IgG1, M1-IgG3, and M1-IgGh systems, respectively. The shades correspond to the standard deviations.

We proceeded to investigate the dynamics of the Fc of the three mAbs relative to M1. We calculated the displacement between the center of mass of the Fc domain of all mAbs and the M1 protein over the time course of the simulations (**Fig. 3A**). The results showed larger displacement for both IgG3 and IgGh with average values of 111.01±24.50 and 167.13±17.60 Å, respectively, compared to IgG1 (94.10±6.77 Å **Supp. Table 2**). The larger standard deviations for both IgG3 and IgGh systems suggest the higher flexibility of the Fc domain in those two systems with respect to the M1 protein. To further analyze the flexibility of the Fc domain in 3D space, we recorded the position of the center of mass of the Fc domain from all three mAbs in every conformation of the MD simulations, with the center of mass of the M1 protein maintained at the origin (**Fig. 3B**). Individual representations for the three systems are depicted in **Figure 3C-3E**. This analysis showed that the Fc domain of both the IgG3 and IgGh systems spans a larger 3D volume compared to the Fc domain of IgG1. Moreover, the traces of the Fc domain from IgG3 showed larger degrees of bending that bring the Fc domain closer to the M1 protein. However, in the case of IgGh, while the Fc domain remained flexible, it kept a certain distance from the M1 protein and explored the space differently. Thus, the hinge of IgG3 leads to both larger degrees of freedom and more flexibility, while the IgG1 hinge confers spatial constraints.

**Figure 3.**
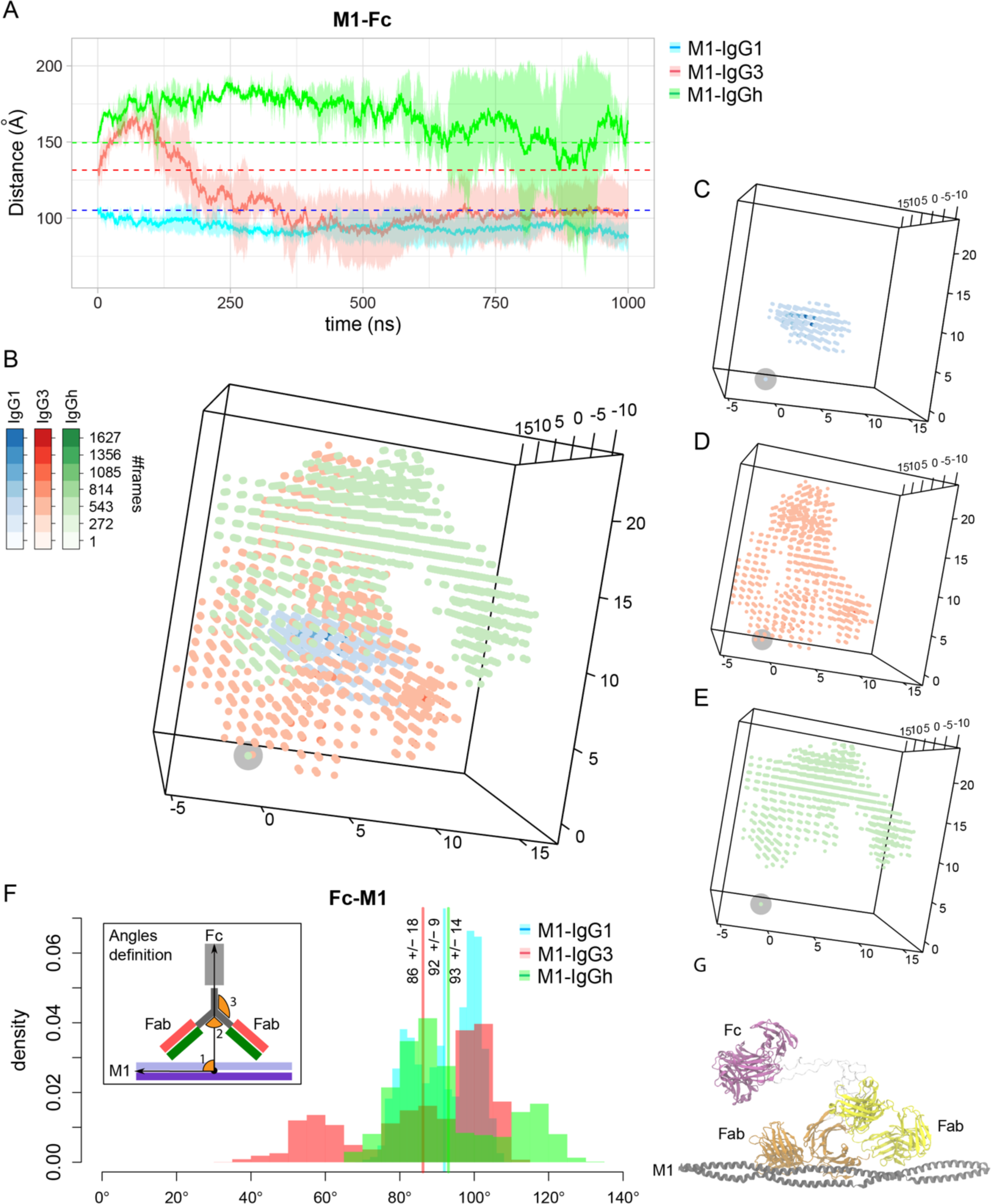
The IgG3 hinge provides broad Fc spatial mobility spectra and flexibility relative to M protein. **A** The average distance between the center of mass of the M1 protein and the Fc domain of the IgG, IgG3, and IgGh was measured over the three replicates of each system (reported in pink and cyan, respectively). The shades correspond to standard deviations, and the initial distances are shown with dashed lines. **B** The center of mass of the Fc domain of IgG1 (in blue), IgG3 (in red), and IgGh (green) at every snapshot of the simulations are depicted. The intensity of the colors represents the population of each point. The center of mass of M1 is shown with a gray circle at the origin. The individual distribution of points are reported for **C** IgG1-M1, **D** IgG3-M1, and **E** IgGh-M1 simulations. The angle changes formed between the Fc domain and protein M1 are reported for IgG1 (in pink), IgG3 (in cyan), and IgGh (green) when both Fabs contact the protein M1. The average values are shown with lines. The vectors forming the angle are depicted in the inset. **G** The most representative conformation of the M1-IgG3 is reported when dual-Fab binding was observed. The IgG-M1 models in the figure are the starting models for the MD simulation.

Furthermore, we measured three sets of angles: *i)* between the Fc domain of the IgGs and the M1 protein, *ii)* between the Fc and Fab domains of the IgGs and *iii)* between the two Fab domains of the IgGs (**Fig. 3F**). These angles were recorded from all MD conformations (**Supp. Fig. 3A-C**); however, only the values where both Fab domains are in the proximity (<30 Å) of the M1 protein are of interest to this study due to the dual-fab binding phenotype of Ab25. Thus, we focused on the conformations where dual-Fab binding occurred (**Supp. Fig. 3 D-F**).

The Fab1-Fab2 angles showed that the antibodies were tilted in different ways but with no substantial variation in the three systems during the simulations **(Supp. Fig. 3F)**. The changes of Fab1-Fab2 angles for IgG1 are in the range of 86° and 139° (with an average value of 111°±15°), for IgG3 between 37° and 80° (with an average value of 62°±8°), and for IgGh in the range of 19° and 57° (with the average value of 38°±7°, **Supp. Table 2**). For the Fc-Fab angles (**Supp. Fig. 3E, Supp. Table 2**), we observed larger distributions of angles for IgG1, IgG3, and IgGh within the ranges of 13° - 114°, 1° - 117°, and 0° - 77°, respectively. The average values of the Fc-Fab angles for the three systems were 66°±22°, 72°±31°, and 31°±14°, respectively. The angle span of the Fc-Fab for IgGh (77°) is roughly two-thirds of the Fc-Fab angle span in IgG1 (101°) and IgG3 (116°) systems. Considering the Fc-M1 angles (Supp. Fig. 2D, Fig. 3F), the following spans were observed: 69° - 108° for IgG1, 34° - 113° for IgG3 and 66° - 132° for IgGh, with the average values of 92° ± 9°, 86° ± 18° and 93° ± 14°, respectively. Thus, IgG3s Fc exhibits twice as large an angle span (79° vs. 39°) and IgGh 1.7-fold (66° vs. 39°) compared to IgG1s Fc relative to the M1 protein (**Supp. Table 2)**.

Models for Fc receptor activation suggest that the accessibility of Fc domains is important to increase the avidity of the interactions (24–25). Therefore, we further analyzed the most prevalent configurations (clusters) regarding the orientation of the Fc relative to the M1 protein for the respective antibodies (**Supp Fig. 4**). Cluster analysis of the MD trajectories revealed that dual-Fab binding occurred in two out of the three most prevalent clusters of the M1-IgG1 system. At the same time, the binding of only Fab2 was observed in the three most significant clusters of the M1-IgG3 system (**Supp. Fig. 4A**). The results showed that, on average, the Fc domain of IgG1 is in a perpendicular-like position with respect to the M1 protein in the three most representative conformations (**Supp. Fig. 4B-D**). The most prevalent configuration for IgG3 Fc is when it bends from the hinge region and fluctuates in parallel to the M1 protein away from the bacterial surface (**Supp. Fig. 4E**, cluster 1 for the IgG3-M1 system **Supp. Fig. 4A**). For IgGh, we detected many clusters with small populations of conformations. This suggests large conformational changes and high flexibility of the IgG hybrid system during the simulations (**Supp Fig. 4A**). These results showed that the IgG3 hinge region increased the movement span. Taken together, the IgG3 hinge region provided IgG3 and IgGh with broad spectra of Fc mobility and flexibility relative to the M1 in 3D space, where IgGh gained IgG3’s increased flexibility through the exchange of IgG1’s hinge with that of IgG3’s.

### The constant domain influences the interaction network between the IgG Fabs and M1 at an atomic level

For the simulations analyzed across all three IgG-M1 systems the binding of Fab2 to the M1 was stable for all three antibodies, while the Fab1 showed a transient behavior and was most stable for IgG1 (**Fig. 2C-D**, **Fig. 2G-H**). One replicate of IgGh became dislodged from M protein during the simulation and was excluded from further analysis. To better understand the binding stability of both Fabs across all MD trajectories, we extracted the network of hydrogen bonds (Hbond) and salt bridges at the interface between the Fab domain of IgGs and the M1 protein (**Fig. 4**). This analysis revealed two stable salt bridges (E44-K58; E46-K65) formed between the VH domain in Fab2 of both IgG1 and IgG3 and the M1 protein (**Fig. 4A-B**). For the IgGh replicates, two salt bridges (E46-K56; E46-K58) occurred, but only one salt bridge per replicate. These salt bridges provide insight into the stability of Fab2 seen during the simulations. Interestingly, although IgGh lacks the presence of a salt bridge in one of the two simulations (in one simulation, the E46-K56 is absent, and in the other, it is E46-K58), the analysis also revealed a new salt bridge between Fab1 and M1 (E76-K127). IgG3 and IgG1 did not share this interaction, and neither of these antibodies formed a salt bridge between their Fab1 and M1.

**Figure 4.**
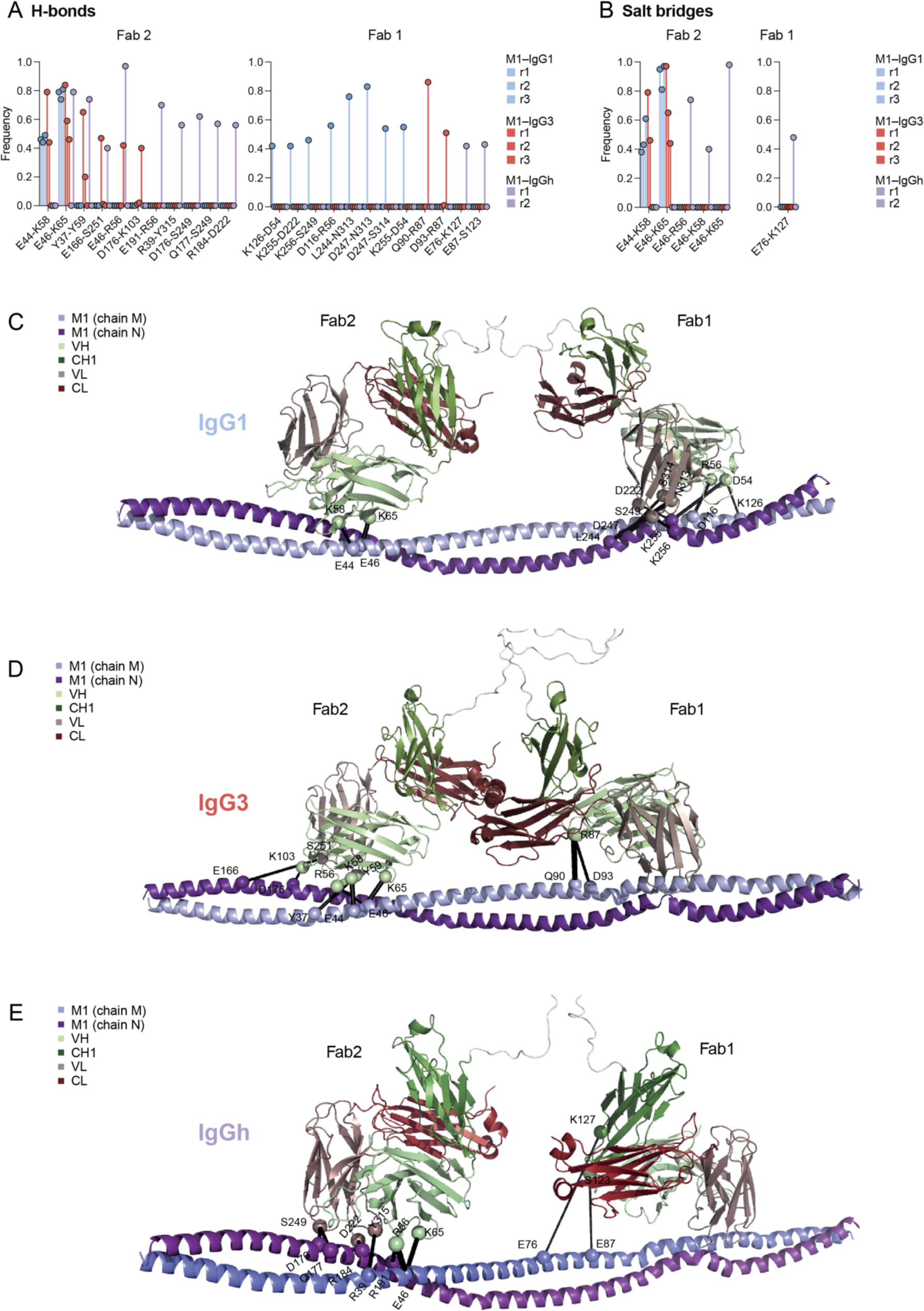
The constant domain influences the interaction network between the IgG Fabs and M1 at an atomic level. The interaction network at the interface of Fab domain and protein M1. The frequency of **A** H-bonds and **B** salt bridges formed in every replicate of the IgG1-M1 and IgG3-M1 simulations is reported in red and blue shades, respectively. The interactions are also shown on the structure of **C** IgG1-M1, **D** IgG3-M1, and **E** IgGh-M1 with black lines. The residues involved in the interactions are shown as spheres. The two chains of protein M1 are colored in light and dark purple, and the VH, CH1, VL, and CL domains are shown in light green, dark green, pink, and red, respectively. The IgG-M1 models in the figure are the starting models for the MD simulations.

Moreover, we recorded a set of eight Hbonds for the Fab1 domain of IgG1, whereas only two Hbonds were observed for the Fab1 domain of IgG3 and IgGh (**Fig. 4C-E**). These hydrogen bonds that IgGh and IgG3 form are not identical and are formed by different amino acids. In addition, these amino acids in Fab1 of IgG3 and IgGh bind to different residues on the M1 protein, which do not interact with Fab1 of IgG1. The reduction of hydrogen bond interactions in Fab1 between IgG3-M1 and IgGh-M1 could explain the less stable binding of Fab1 during the simulations. Furthermore, there are notable differences between the three IgG-M1 systems in the Fab2 hydrogen bond interaction network. IgG3 and IgGh Fab2 both form six and seven Hbonds, respectively, with the M1 epitope, while IgG1 Fab2 forms only two (**Fig. 4C-E**). It is important to note that the Fab2-M1 interactions for all three antibodies are very stable in the simulation (**Fig. 2C-D**, **Fig. 2G-H**). IgG1s Fab2, despite the reduced amount of Hbonds, exhibits stable binding to M1, most likely due to the presence of two salt bridges mentioned prior (E44-K58; E46-K65). The two Hbonds of IgG1 in the Fab2 (E44-K58; E46-K65) are both shared by IgG3, and IgGh shares one (E46-K65). One of the remaining Hbonds is shared between the IgGh and IgG3 Fab2 (R46-R56). Different amino acids form the remaining Hbonds. Taken together, the analysis of the network of interactions reveals differences in both Fabs across all three IgG-M1 systems. The differences between IgG3 and IgG1 in Fab1 could explain the reduced binding of IgG3 to M1 measured previously (**Fig. 1B**). The Fab2 interactions with M1 appears more stable due to the salt bridges in all IgG systems. In terms of hydrogen bond formation, the IgGh network of interactions seems to be similar to IgG3. However, an exception is that IgGh Fab1 forms a new salt bridge with M1. Thus, despite the reduction of Hbonds in IgGh Fab1-M1, IgGh binding to M1 is compensated by the presence of this novel salt bridge (E76-K127) which could explain its IgG1-like reactivity to M1. Nonetheless, our analysis revealed that the network of atomic interactions between the antigen and antibody Fabs is altered when exchanging parts of (IgG3 hinge region) or the entire constant domain.

### IgGh retains the high affinity of IgG1 and gains superior opsonic function

The MD simulations highlighted the impact of the IgG3 hinge on increased spatial Fc movement and relative flexibility to the M1 protein. To investigate how the Fc mobility and flexibility influence antibody function, we compared the IgGh opsonic function to that of its parent subclasses (**Fig. 5A**). First, we analyzed IgGh’s ability to promote phagocyte-bacteria association by calculating the MOP_50_. In these sets of experiments, the differences in MOP_50_ between IgG3 and IgG1 were 3-fold (IgG3=21, and IgG1=62) (**Fig. 5B-C**). Adding the IgG3 hinge improved the opsonic efficacy for IgGh 2.5-fold over IgG1 (MOP_50_= 25, **Fig. 5B-C**). All three mAbs had a much higher association than the negative control (MOP_50_= 188), also reflected in the individual MOP_50_ analysis (**Fig. 5D**). We further assessed the importance of hinge flexibility by looking at the proportion of THP-1 cells with internalized bacteria and the bacterial fluorescence. The IgGh outperformed both parent subclasses in internalization with more phagocytes internalizing bacteria (**Fig. 5E**). We also analyzed the quantity of bacteria phagocytosed, which was the highest with IgGh as the opsonin (**Fig. 5F**). Finally, we calculated a phagocytosis score (percentage of associating cells of the whole population multiplied by the bacterial signal of the whole cell population), and the results from this analysis showed that IgGh exceeds both the potent IgG3 and the original IgG1 (**Fig. 5G-H**). At MOP 200, IgGh has a 2.5-fold higher phagocytosis score than IgG3 (50.400 vs 20.600), 4-fold over IgG1 (12900), and 35-fold over control (1400). Taken together, these results strengthen the hypothesis that the spatial Fc mobility and flexibility enabled by IgG3’s hinge is a crucial factor for potentiating Fc-mediated phagocytosis. Our data also showed that IgGh is a more potent opsonin than IgG3 when all four metrics are considered together.

**Figure 5.**
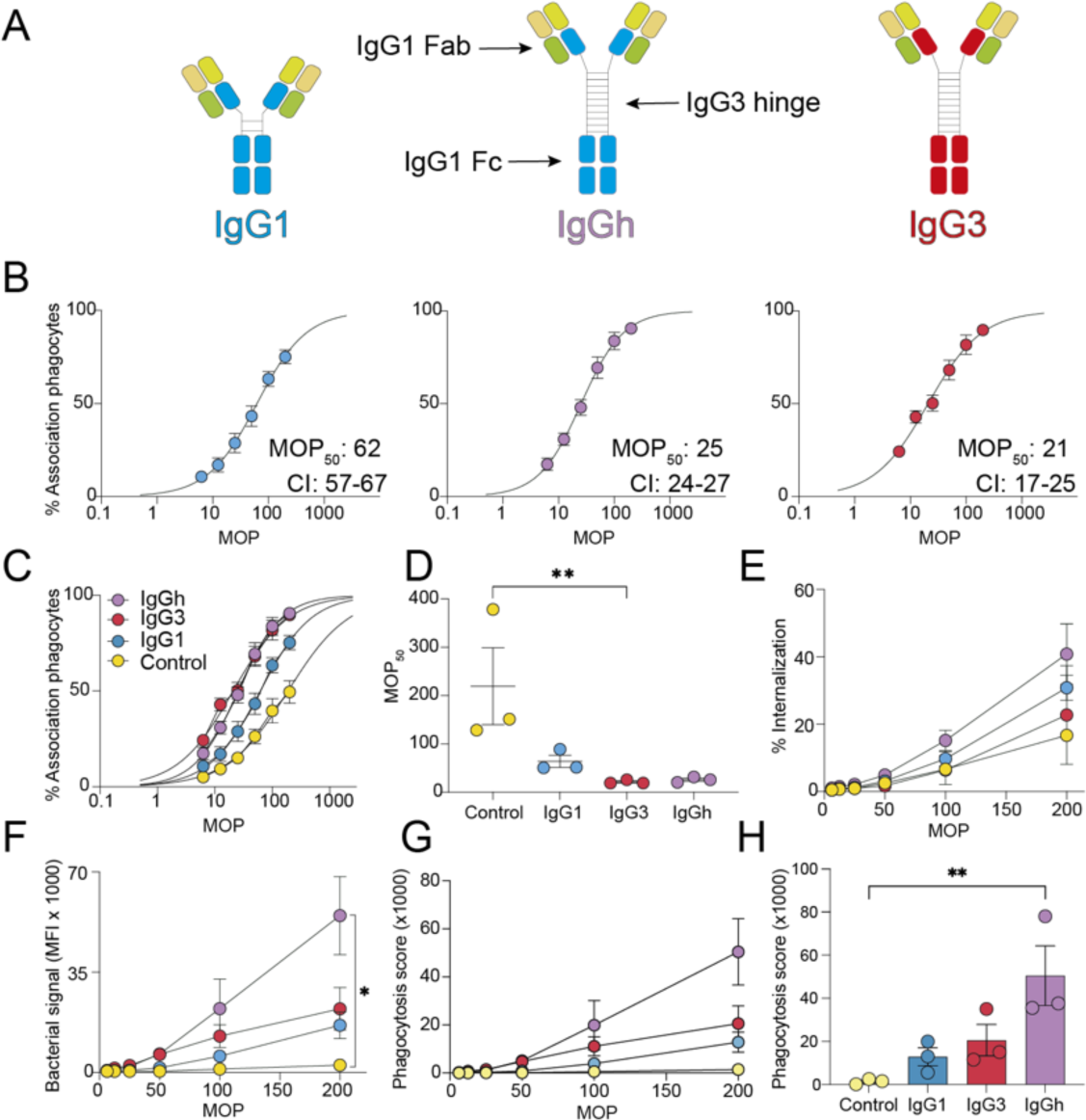
IgGh retains the high affinity of IgG1 and gains superior opsonic function. **A** Schematic of IgG1, IgG3, and the IgGh The antibodies consist of two identical heavy chains and identical light chains. The variable heavy domain (yellow), the constant domain of the light chain (dark green), and the variable light domain (light green) are identical for the three antibodies. The constant domains of the heavy chains are depicted in different colors: blue for IgG1 and red for IgG3. **B** MOP_50_ curves of the subclasses, bacteria were opsonized with 15 ug/mL of each respective antibody. The Y-axis shows the percentage of THP-1 cells that are positive for bacteria, while the X-axis depicts the ratio of bacteria to phagocytes added (MOP). The MOP_50_ values with a 95% confidence interval are shown in each graph. The data points are from three independent experiments. **C** Shows the MOP_50_ curves of all the antibody treatments in one graph. Control is Xolair, human IgG1 anti-IgE. **D** Individual MOP_50_ values from the three independent experiments. **E** Percentage of THP-1 cells with internalized bacteria across the MOP range. **F** Amount of bacteria that are phagocytosed by the whole THP-1 cell population measured as median MFI of FITC-A (bacterial signal, Oregon Green). **G-H** Phagocytosis score for each antibody treatment is shown with MOP on the X-axis for **G**, while in **H**, the MOP is 200. All statistical analysis was done, comparing the treatments to the control antibody with Kruskal-Wallis multiple comparisons corrected by Dunn’s correction test. * denotes P-value < 0.05 and P-value > 0.05 is ns. The data points in figure **B-I** represent the mean value, and the error bars are in SEM.

### Increased hinge flexibility of IgG1 confers antibody protection against systemic infection in mice

To study the biological relevance of increased Fc flexibility, we proceeded with in vivo experiments. We used an animal model of invasive infection and passive immunization where C57BL/6J female mice were prophylactically pretreated 6 hours before infection with IgG1, IgG3, IgGh, or PBS (Fig. 6A). Bacteria were then administered on the flank in a low volume to generate a localized tissue infection, which differs from the previous model used to assess Ab25 IgG1, where the mice were injected with higher volumes in the scruff to generate a diffuse tissue infection (11). We assessed the protective effect of the antibodies by quantifying bacterial dissemination to distant organs (spleen, liver, and kidneys) and systemic cytokine mobilization in blood.

**Figure 6.**
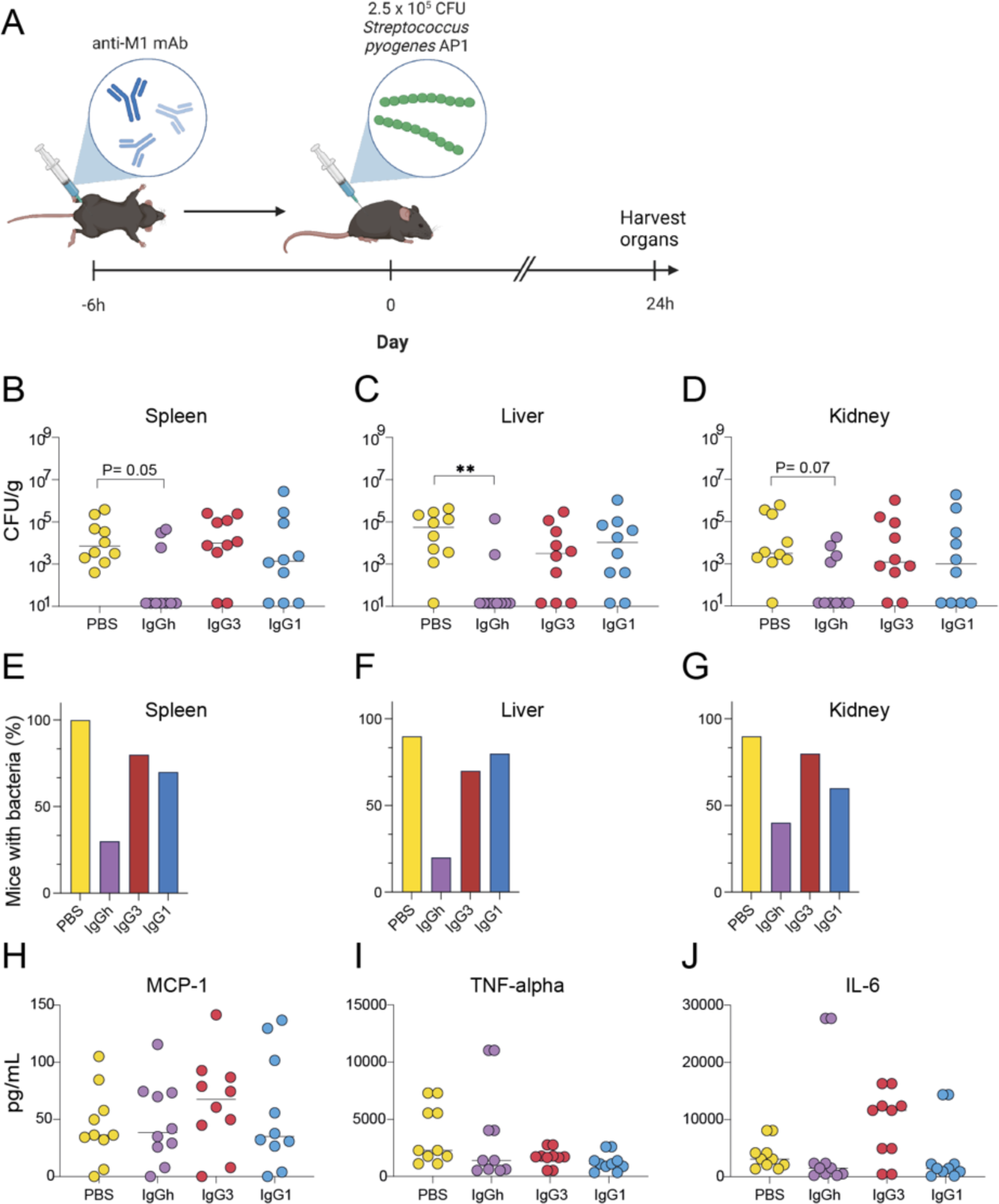
Increased hinge flexibility of IgG1 confers antibody protection against systemic infection in mice. **A** Schematic of the animal experiment: pre-treatment 0.4 mg of antibody treatment or PBS 6 hours before infection with AP1. Animals were sacrificed 24 hours post-infection, and organs and blood were harvested for analysis. **B-D** Shows pooled data from two independent experiments of 5 mice in each cohort. Bacterial load (CFU/ g) in each organ was determined by serial dilution and viable count determination after overnight incubation. **E-G** shows the percentage of mice positive for bacteria in each respective organ for all treatments. **H-J** Cytokine levels (pg/ ml) in plasma were measured using a cytometric bead array. In **B-C**, the median is shown. Statistical analysis was performed, comparing the treatments to the PBS, by Kruskal-Wallis with multiple comparisons and Dunn’s correction test. ** denotes P-value < 0.01, * denotes P< 0.05 and P-value > 0.05 is ns.

Interestingly, in this model, both IgG3 Ab25 and IgG1 Ab25 treatment of mice reduced bacterial spread to distant organs compared to the PBS control but did not completely prevent systemic dissemination. Distinct organotypic patterns of dissemination were observed (**Fig. 6B-D**). IgG3-treated mice did not reduce bacterial load in the spleen compared to PBS (median 9800 CFU/g vs. 7200, **Fig. 6B**), while mice treated with IgG1 showed a reduced spread to the spleen (median 1400 CFU/g **Fig. 6B**). In the liver, IgG3 reduced the bacterial load to a greater extent than IgG1 (median 3200 CFU/g vs. 10800) and compared to the PBS group (median 55600 CFU/g, **Fig. 6C**). In the kidneys, there was a decrease in bacteria for mice in both IgG1 and IgG3-treated groups (median 1000 CFU/g and 1200 respectively) compared to the PBS-treated mice (median 3200 CFU/g, **Fig. 6D**). In contrast, IgGh did not have any detectable bacterial presence for a majority of the cohort (undetectable bacteria were labeled as 14 CFU/g in the graph, IgGh had median 14 CFU/g for all organs, **Fig. 6B-D**). More importantly, the control group only had one mouse (10%) in the cohort that was bacteria-free in all three organs (**Fig. 6E-F**). For IgG1, 30% of all mice were bacteria-free in the spleen (**Fig. 6E**), 20% in the liver (**Fig. 6F**), and 40% in the kidneys (**Fig. 6G**). This was comparable to the IgG3 response, with 20% for the spleen, 30% for the liver, and 20% for the kidneys. IgGh stands out in this analysis, with 70% of the cohort not having bacteria in the spleen, 80% in the liver, and 60% in the kidneys (**Fig. 6E-G**).

We analyzed cytokine expression in the blood of each respective mouse in all groups. Large individual variation in each group was observed for MCP-1, TNF-alpha, and IL-6. All three mAb-treated groups had equal MCP-1 compared to the control group (**Fig. 6H**) and similar TNF-alpha response (**Fig. 6I**). For IL-6, there was a clear distinction between IgG3-treated groups and the other groups (**Fig. 6J**). The almost complete prevention of bacterial dissemination to distant organs combined shows that the IgGh is the most protective antibody. Although IgG3 has extensive Fc-mobility equal to IgGh, it did not provide similar levels of protection, possibly explained by its reduced half-life, lower affinity, or different Fc-receptor affinity towards mouse Fc-gamma receptors. However, comparing IgG1 and IgGh does not suffer from these confounders. It highlights that Fc spatial flexibility and mobility can be determining factors for complete immune protection against systemic streptococcal infection.

## Discussion

Due to significant technological developments with next-generation sequencing, single B-cell technology, and phage display, the generation of clinically relevant and targeted therapeutics antibodies has become more efficient (26). Antibody-based therapeutics are emerging as a rapidly growing class of drugs with a great span in disease targets, ranging from oncology to rheumatology (27). In the field of infectious diseases, clinical success has been chiefly made with antibodies targeting viral pathogens such as RSV (28), HIV (29), and SARS-CoV-2 (30). Today, most antibodies considered for clinical use are Fc-engineered to exhibit beneficial pharmacokinetics, such as longer half-life (27, 28). However, few clinically approved mAbs have been modified to include beneficial properties of the different subclasses, such as inserting the IgG3 hinge in the backbone of IgG1 to increase favorable pharmacodynamic traits.

Pre-clinical studies have shown that subclass-switching to IgG3 enhances functional output *in vitro.* Recently, it was demonstrated that neutralization was potentiated when subclass-switching from an IgG1 anti-Sars-CoV-2 to an IgG3 mAb (31). We have also recently shown that anti-spike SARS-CoV-2 IgG3 mAbs exhibit more potent Fc-mediated functions compared to IgG1 (32). Several other examples exist where IgG3-switched mAbs can exhibit more efficient phagocytosis than IgG1. These cases include antibodies against a variety of pathogens ranging from HIV-1 (23), staphylococci (21), meningococci (33), to pneumococci (22). Despite the promise of the IgG3 subclass, its use as a template has been hindered due to a lower half-life in serum (1) and a tendency to aggregate when produced on a large scale (34). Through accomplishments with Fc-engineering, these issues have been remedied (22, 35), making this subclass more appealing for clinical application.

In this work, we generated and characterized all four human subclasses of Ab25, in which we observed an exciting phenotype in the IgG3 subclass. It exhibited reduced binding but improved function. We further investigated this reduced reactivity of IgG3 (and the possible impact of hinge-engineering with IgGh) by looking at the Hbonds and salt bridges of the M1-IgG interaction. Our analysis of the M1-IgG3 interactions revealed a possible mechanism for the reduced reactivity towards M1. Exchanging constant domains from IgG1 to that of IgG3 induced rearrangement of the hydrogen bond network between the Fab domains and the M1 protein. In the case of IgG1, the Hbond network is mainly formed between the Fab1 domain and the M1 protein. However, this network is shifted toward Fab2 and M1 protein in the case of IgG3. While a similar behavior was observed for IgGh, a novel salt bridge in the Fab1 seems to compensate for Hbond rearrangement and maintain an IgG1-like reactivity to M1. While our results agree with other emerging findings on the influence of the constant domain on the binding properties of mAbs, we reveal the details of the loss of binding at an atomic level. We extend the established work of others (4–7) by providing a more detailed mechanism for the loss of binding observed with our *in silico* data. The growing body of evidence showing that the constant domain and variable domain are dependent on each other is challenging the traditional model of variable and constant domain independence (4–7).

The enhanced performance of the IgGh compared to IgG1 shows the functional potency of the IgG3 hinge region. Similar work has been done recently with antibodies against adenovirus and HIV (23,36). Both studies hypothesized that the functional output associated with IgG3’s hinge is due to increased flexibility but did not pursue this hypothesis. We build on those studies and, using *in silico* analysis, reveal that both the IgGh and IgG3 antibodies possess an increased flexibility of Fc due to their extended hinge region. Although the subject of IgG subclass flexibility has been investigated previously (8), our findings highlight molecular differences between IgG1 and IgG3. In the aforementioned study (8), hinge flexibility, measured as SD of Fab-Fc angle, was reported to be ±36° for IgG3 and ±30° for IgG1, with minor differences between the two and no difference in the total Fab-Fc angle span. It is worth noting that these experiments were performed in the absence of the antigen and only entailed the inherent Fc flexibility relative to the antibody itself (Fc contra Fabs).

Our investigations replicate the findings that IgG3 has a slightly larger Fab-Fc angle span (116°±31°) and SD than IgG1 (101°±22°). However, the Fab-Fc angle span for IgG3 also exceeds that of IgGh (77°±14), which rather speaks against the fact that the IgG3 hinge would lead to a higher Fab-Fc angle. More strikingly, we observed much greater flexibility regarding absolute Fc movement, spatial movement in 3D space, and Fc-antigen angle span for the IgG3 and IgGh antibodies which is more of a functional flexibility. To our knowledge, this unique behavior associated with IgG3’s hinge has not been reported before. The differences in Fc flexibility described in this study are much larger than previously thought. We believe this is because we studied the functional Fc flexibility (relative to antigen) and not the inherent flexibility relative to the antibody Fabs and that we could use dynamic metrics (absolute movement, angle of movement, and movement of Fc in 3D space). The functional impact of antibody Fc flexibility shown in this work motivates the necessity of studying these interactions with other antigens. MD is an excellent tool for studying dynamic processes (including protein flexibility) as opposed to analyzing snapshot moments, such as the models determined by single-particle Cryo-EM or X-ray crystallography. We propose that future work on subjects concerning antibody-antigen biophysical interaction (such as flexibility) would greatly benefit from using MD simulations as done here to study functional flexibility and other dynamic antibody interactions. We believe the findings described here have significant implications for our understanding of how antibodies mediate their function and warrant future investigation with other antibodies binding to other antigens.

Our *in vitro* results show that the IgG3 hinge increases opsonic ability. We believe this is a result of increased Fc to Fc-receptor interactions. We do not believe this increased efficacy is due to altered affinity to Fc-receptors. Instead, we theorize that the increased flexibility that the IgG3’s hinge grants enables a higher probability of interacting with Fc-receptors. This increased interaction would enable more IgG3 and IgGh antibodies to interact with Fc-receptors than the IgG1 antibodies. This rationale aligns with models proposed for Fc-receptor activation and initiation of phagocytosis, especially the avidity- and Fc-clustering model (37). Fc-FcR interactions have also been shown to be increased when antibodies bind with low affinity to their targets (38). This could be an alternative explanation for IgG3’s opsonic performance but would not explain IgGh’s efficacy since it has a higher affinity towards the M1 protein.

We provide *in vivo* pharmacodynamics of hinge engineering, complementing previous pharmacokinetics (biostability in serum) studies (23). Our work shows that utilizing IgG3’s flexible hinge, and thus increasing antibody flexibility, has direct *in vivo* implications. IgGh protects against systemic bacterial dissemination, while the more rigid IgG1 does not. It is worth noting that IgG1 Ab25 showed a more prominent *in vivo* effect in a previously used animal model (11), while in the current model, although more protective than the control, the result was not as strong as we observed in the previous model. The main difference between the models lies in the route of administration that generates a diffuse (scruff) or restricted (flank) local infection. The route of administration can affect the host-pathogen interactions underlying invasion and disease progression, but understanding the mechanisms involved would require further investigation. Moreover, the differences between IgGh and IgG3 are worth discussing. Both mAbs have similar flexibility and opsonic efficacy, but the IgGh has a higher affinity towards the M1 protein. In addition, human IgG1 has a higher half-life in mice than human IgG3 (5 vs. 1 day, respectively) (39). A lower half-life and lower affinity to the M1 protein could lead to less IgG3 being functionally active and present during the experiment than IgGh and IgG1. Additionally, mouse phagocytes have different expression of Fc-receptors compared to human phagocytes (40). These Fc-receptors have similar affinities as their human counterparts to human subclasses (41). Despite this apparent conserved binding, human IgG3 antibodies have been shown to induce weaker effector responses by mouse phagocytes (42–43). This is contrary to work with human phagocytes, where IgG3 has been shown to be more potent in inducing effector functions (21–23,33). Thus, the *in vivo* experiments should be interpreted cautiously when comparing IgG3 with that of both IgG1 and IgGh. However, since the IgGh shares the same sequences recognizing Fc-R as IgG1, the results of the *in vivo* experiments serve as a strong proof of concept regarding the clinical application of modifying antibody hinge flexibility. Our study, combined with previous findings (23,36), shows the benefit of Fc-engineering an IgG3 hinge into an IgG1 backbone to enhance the Fc-effector functions of promising mAb candidates. Finally, in the context of mAbs against *S. pyogenes*, our results suggest that the engineered IgGh Ab25 could be an even more promising candidate than IgG1 Ab25 (11) for treating severe streptococcal infection.

## Materials and Methods

### Plasmid generation, transformation, and plasmid purification

To generate heavy chain plasmids of the different subclasses, pVITRO1-Trastuzumab-IgG2 (Addgene plasmid #61884; RRID: Addgene_61884), IgG3 (Addgene plasmid #61885; RRID:Addgene_61885), and IgG4 (Addgene plasmid #61887; RRID:Addgene_61887) was a gift from Andrew Beavils lab (44). The heavy chain and light chain of Ab25 were generated as described previously in Bahnan et al (11). The cloning was done by amplifying the constant region of the heavy chains for IgG2, 3, and 4 by using the high-fidelity Phusion (NEB Biolabs) 3-step PCR protocol. The original heavy chain of Ab25 was linearized so that the constant region of IgG1 was removed. The primer design was planned so the amplified vector and insert contain overlapping ends, which are compatible with HIFI DNA assembly (NEB Biolabs). For the generation of the IgG1/3 hybrid, a 2-step PCR was done to amplify the insert and the vector, and the fragments were combined by HIFI DNA assembly. The plasmids were transformed into competent *E. coli* supplied by the manufacturer of the HIFI-DNA assembly according to their instructions.

Following transformation, single colonies were picked and grown for 16 h 3 mL Luria Bertani (LB) medium containing 10 μg/mL ampicillin in a shaking incubator at 37 °C. The following day the bacteria culture was harvested and used for miniprep (QIAprep Spin Miniprep Kit, QIAGEN) to acquire pure plasmids. The plasmid concentration was measured by drop spectrometry (DeNovix) and was sent for sequencing to verify that no mutations had been introduced.

For midi-preps, LB was inoculated with a single colony in 3 mL LB containing 10 μg/mL ampicillin and grown for 6 h in a shaking incubator at 37 °C. They were added to 27 mL LB containing 10 μg/mL ampicillin for 16 h under the same conditions. The culture was harvested the day after, and the plasmids were purified by QIAGEN plasmid plus midi kit according to the manufacturer’s instructions.

### Bacteria culturing

*S. pyogenes* SF370 (*emm1* type) was streaked on a plate with Todd-Hewitt Yeast (THY) agar with erythromycin (1:1000 dilution, 100 mg/mL) when the strain containing the GFP plasmid was grown. When the WT SF370 strain was grown, this was without erythromycin. The plate was kept in use for up to 2 weeks. The day before an experiment, one colony was added to 10 mL of THY media with erythromycin for the GFP expressing SF370 and without for the WT. After 16 hours, the overnight was diluted 1:20 and grown to mid-log for 2.5 h for the GFP and 2hr for the wild type.

For the phagocytosis assay, the bacteria were heat-killed. After growing to mid-log, the cultures were washed once with PBS, put on ice for 15 minutes, put in a shaking heat block (300 rpm) at 80 °C for 5 minutes, and then directly put back on ice for 15 minutes. The heat-killed bacteria were stained 1:500 with Oregon Green 488-X succinimidyl ester (2 μL of dye from stock in 1 mL of PBS) (Thermo Fisher) fluorescent dye for 30 minutes at 37 °C. After staining, the bacteria were washed with PBS, resuspended with sodium carbonate buffer (0.1M pH 8.3), and stained with CypHer5E (Fisher Scientific) 1:100 in a volume of 200 µL for 30 minutes at 37 °C, protected from light. The bacteria were washed twice with PBS and then resuspended in sodium medium (5.6 mM glucose, 127 mM NaCl, 10.8 mM KCl, 2.4 mM KH_2_PO_4_, 1.6 mM MgSO_4_, 10 mM HEPES, 1.8 mM CaCl_2_; pH adjusted to 7.3 with NaOH) and stored at 4 °C.

### Cell culture, antibody production, and antibody purification

THP-1 monocytic cells were cultured in RPMI with 10% FBS and L-glutamine and were kept at 5-10×10^5^ cells per mL. The cells were never used more than 6 weeks after thawing the vials.

6.5 million human embryonic kidney (HEK) 293 cells (Gibco) were seeded the day before transfection in 150mm dishes in DMEM (Gibco) supplemented with 10% FBS (Gibco) and L-glutamine. Transfection was done by first treating the cells with chloroquine diluted 1:1000 in fresh DMEM to replace the old. 6 h after Chloroquine treatment, 20 µg of Heavy chain and Light chain plasmid was diluted in 1mL OptiMEM(Gibco) and 120 µL polyethyleneimine (1 mg/mL PEI, Thermo fisher) for each plate. The mix was left to incubate for 20 minutes at room temperature before gently being added to the plates. The following day the DMEM was replaced by OptiMEM with 2 wash steps with sterile PBS before the switch. The cells were left to incubate for 48 h, and the supernatants were collected and spun for 5 minutes at 450 x g to remove any cells.

Expi293F suspension cells were purchased from Gibco (Thermo Fisher) and routinely cultured in 125 mL Erlenmeyer flasks (Nalgene) in 30 mL Expi293 medium (Gibco) in an Eppendorf s41i shaker incubator at 37°C, 8% CO_2_, 120 rpm. Cells were passaged and split to a density of 0.5 x 10^6^ cells/mL every 3 to 4 days. The day before transfection, the cells were seeded at a density of 2 x 10^6^ cells/mL. The next day, cells were seeded at 7.5 x 10^7^ cells in 25.5 mL Expi293 medium. 20 μg of respective heavy and light chain plasmids were used to transfect Expi293 cells. Expi293 cells were transfected at a concentration of 3 million/mL in a volume of 25.5mL in Expi293 media. The plasmids were mixed with 2.9 mL of OptiMEM media and 100 µL of Expifectamine. The mix was left to incubate at room temperature for 15 min. After incubation, the mix was added to the cell flasks containing the cells. 22 hours later, 1.5 mL of enhancer 1 and 0.15 mL of enhancer 2 (Expifectamine kit, Gibco) were added to the cells. 72 hours later, the cell’s supernatant was harvested.

The supernatants (from both methods) were later purified the same day by incubation for 2 h with protein G sepharose (Cytiva). After incubation, the beads were washed 2 times, 50 mL PBS in a chromatography column (Biorad). Elution was done by adding 2×5 mL of HCl-glycine (0.1M, pH 2.5), and 1000 mL of TRIS (pH 8, 1M) was used to neutralize the pH. The purified antibodies were concentrated, and the buffer was exchanged to PBS by means of spin columns. The antibody concentrations were measured by doing an SDS-PAGE with serial dilutions of Xolair and Denovix drop spectroscopy using the IgG setting.

### Binding assays with secondary conjugated antibodies

For binding assays, live *S.pyogenes* strain SF370 transformed with GFP was used. 10mL of mid-log culture was concentrated and resuspended in a final volume of 1mL in PBS. The bacteria were sonicated for 2×2 minutes to break clumps. 10 µL of concentrated bacteria were added to each well of a 96-well low-binding plate containing 90 µL of anti-M antibodies (concentrations of antibodies varying between 20-0.65 µg/mL) in PBS. Serial dilutions of antibodies were prepared with concentrations ranging from 20 µg/mL to 0.65 µg/mL. The bacteria were opsonized for 30 min at 37 °C with gentle shaking. After opsonization, the bacteria were washed twice with PBS before adding 50 µL (1:500 dilution from stock) secondary antibody, Alexa Fluor® 647 AffiniPure F(ab’)₂ Fragment Goat Anti-Human IgG, F(ab’)₂ fragment specific, (109-606-097, Jackson Immunoresearch). The secondary antibody was incubated for 30 minutes at 37 °C with gentle shaking. Following this, PBS was added so that the final volume reached 250 µL. The plates were analyzed by using a Cytoflex flow cytometer (Beckman Coulter). The data acquired from the flow cytometer experiments described above were processed on Flowjo software for gating strategy. The positive gate for the live GFP-expressing bacteria was set by gating for size (SSC) and GFP fluorescence (FITC) (**Supp Fig. 1A**). The gate for antibody reactivity to M was set by using a low concentration of Xolair (0.5 micrograms per mL) for Fc-background binding (**Supp. Fig. 1A**). Antibody reactivity was determined by gating on percentage bound bacteria positive for the secondary antibody (APC) **(Supp. Fig. 1A)**. The data were analyzed in GraphPad Prism, and the percentage of parents was plotted against antibody concentration followed by a non-linear regression analysis with Bmax constrained to 100% and K_D_ unconstrained.

### Directly conjugated antibody binding assay

The antibodies were conjugated with Alexa Flour 647 using the GlyClick kit (Genovis) according to the manufacturer’s protocol. The binding assay was performed with live SF370 expressing GFP. 10 mL of mid-log of culture was spun down and taken up in 1 mL PBS (10X concentration). The bacteria were sonicated as described previously. 10 μL of the bacteria from the 10x stock was added to 90 μL of antibodies in PBS. The concentrations of antibodies varied from 10-0.6 μg/mL for IgG1 and the hybrid. The bacteria were opsonized for 30 min at 37 °C with gentle shaking. 150 μL of PBS was added, and the plate was washed once and resuspended in a final volume of 150 μL. The plates were analyzed by using a Cytoflex flow cytometer (Beckman Coulter). The data acquired from the flow cytometer experiments described above were processed on Flowjo software for gating strategy. The positive gate for the live GFP-expressing bacteria was set by gating for granularity (SSC) and GFP fluorescence (FITC) (**Supp. Fig 2A**). Antibody reactivity was determined by gating on percentage bound bacteria positive for APC-A (**Supp. Fig 2A**). The data were analyzed in GraphPad Prism, and the percentage of bacteria positive for antibody was plotted on the Y-axis with antibody concentration on the X-axis. A non-linear regression analysis was performed, with Bmax constraint set to 100% while Hill-slope and K_D_ were unconstrained.

### Phagocytosis assay

After heat killing and staining the bacteria as described above, the samples were sonicated for 4 minutes to remove big clumps of aggregated bacteria. The bacteria were diluted 1:500, and 250 µL were added to a 96-well plate and run on Cytoflex flow cytometer (Beckman Coulter) to quantify the concentration as events/µL. In addition, to control the pH responsiveness of the CypHer5e staining, 2 µL of sodium acetate (3M, pH 5.0) was added to the wells to observe a right shift in the APC channel, indicating successful staining.

Serial dilutions of bacteria were done with MOP (bacteria per phagocyte) as the variable, and each well was opsonized with 15 µg/mL of anti-M antibodies or Fc-control for 45 minutes at 37 °C gently shaking. 100 µL was the end volume before the addition of the cells. While the plate was opsonizing, the THP-1 cells were collected and washed once to remove RPMI and exchanged to sodium media. THP-1 cells, numbering 1 x 10^5^ cells, were added at a concentration of 2 x 10^6^ per mL after being on ice for 5 minutes. The cells were left phagocytosing for 30 minutes at 37°C while gently shaking. The plate was put on ice for 15 minutes to stop further phagocytosis. The plate was analyzed in the Cytoflex flow cytometer (Beckman Coulter) with 488nm and 638nm lasers and filters 525/40 and 660/10 APC. For bacteria, the threshold was set to FSC-H 2000, SSC-H 2000, and for THP-1 cells, FSC-H 70000. The gain was set to 20 for FITC and 265 for APC. The acquisition was set to capture 5000 THP-1 cells with medium velocity (30 µL/min). While running, the 96-well plate was kept on an ice-cold insert. THP-1 cells were gated first by size (FSC) and granularity (SSC) and then for single cells by SSC-A and SSC-H. By using untreated THP-1 cells, we set a gate for non-associated THP-1 cells, associated with bacteria and associated with bacteria and internalized by using the FITC and APC signal (**Supp. Fig 1B** and **Supp Fig 5B**). The data acquired from the flow cytometer experiments described above were processed on Flowjo software for gating strategy and later analyzed on Graphpad Prism. The MOP_50_ analysis was calculated by using the PAN-assay model as described in Prism (20), which in short, is a non-linear regression model where EC_50_ of MOP is calculated based on the persistent association (association) as an outcome metric with a top value set to 100% and bottom value constrained to 0%. Finally, the phagocytosis score was calculated by multiplying the percentage of bacteria-positive THP-1 cells (associating) with the bacterial signal (MFI, Oregon green) for the whole population of THP-1 cells.

### Animal model

All animal use and procedures were approved by the local Malmö/Lund Institutional Animal Care and Use Committee, ethical permit number 03681-2019. Nine-week-old female C57BL/6J mice (Scanbur/ Charles River Laboratories) were used. Monoclonal antibodies (0.4 mg/mouse) were administered intraperitoneally 6 h pre-infection. The pretreatment groups were coded, and the experimenters were blinded to which group had which intervention, and unmasking was done after the data had been analyzed and compiled. *S. pyogenes* AP1 was grown to an exponential phase in Todd–Hewitt broth (37°C, 5% CO^2^). Bacteria were washed and resuspended in sterile PBS. 2.5 x10^5^ CFU of bacteria were injected subcutaneously on the right flank, leading to a local infection that progressed to systemic infection within 24 h. Mice were sacrificed 24 h post-infection, and organs (blood, livers, spleens, and kidneys) were harvested to determine the degree of bacterial dissemination. Cytokines in plasma were quantified using a cytometric bead assay (CBA mouse inflammation kit, BD biosciences) according to manufacturer instructions. The blood from the mice was analyzed by Western blots to control for human IgG integrity at the end of the experiment. We found that most IgG remained intact, and thus appeared unaffected by potential hydrolysis by bacterial enzymes or other potential degradation.

### Structural models

The Fc and Fab domains of IgG1 and IgG3 were *de novo* modeled separately, using AlphaFold2 (45–46), considering the homo-oligomer state of 1:1. The multiple sequence alignments were generated by MMseqs2 (47). The output models were then relaxed using Rosetta relax protocol (48), where the side chains and disulfide bridges were adjusted. For IgG3, as the hinge region is long and contains multiple disulfide bridges, it was modeled separately using AlphaFold2 and relaxed by Rosetta relax protocol. The loops connecting the hinge region of both IgGs to Fc and Fab domains were then re-modeled and characterized using the DaReUS-Loop web server (49–50). Finally, the full-length structure of both IgGs was relaxed, and all disulfide bridges (specifically in the hinge region) were adjusted using Rosetta relax protocol. The M1-IgG models were generated using a targeted cross-linking mass spectrometry (TX-MS) approach (51). The cross-link constraints (XLs) were derived from a set of experiments generated in our previous study (11), and the threshold of 32 Å was applied to map XLs on the structure. Accordingly, two epitopes of the M1 protein in the B3 and C1 domains interact with the Fab domains of IgGs. The M1-Fab interactions for both IgGs were also separately characterized and re-adjusted using AlphaFold before relaxing the final structure using Rosetta relax protocol.

### Molecular dynamics simulations

We performed three replicates of 1 µs MD simulations starting from the models obtained for M1-IgG1, M1-IgG3, and M1-IgGh systems. This resulted in a total of 9 µs MD simulations. In one replicate of the M1-IgGh system, we observed large conformational changes in the IgGh with respect to the M1. In this replicate, the Fab domains were detached, resulting in a global shift of the antibody and, finally, attachment of the Fc domain to the M1. Therefore, this replicate was discarded from the analysis. The 3D coordinates of the two models generated for M1-IgG1 and M1-IgG3, described in the previous section, were retrieved. Both IgG1 and IgG3 have 6 chains (A-F); four disulfide bonds are present on the Fc domain, and four on each Fab domain. However, the IgG1 has a shorter hinge region of 14 residues (E222-P236) with 2 disulfide bonds, while the hinge region of the IgG3 is composed of 61 residues (E222-P283) and 11 disulfide bonds. The M1 protein has 2 chains (M, N), and for the MD simulations, we considered its central region (residues A149-L291 on chains M and N). MD simulations were carried out with the GROMACS 2020.6 (52) using the CHARMM36m force field parameter set (48): (i) Na^+^ and Cl^-^ counter-ions were added to reproduce physiological salt concentration (150 mM solution of sodium chloride), (ii) the solute was hydrated with a triclinic box of explicit TIP3P water molecules with a buffering distance of up to 12 Å, and (iii) the environment of the histidine was checked using MolProbity (53). First, we performed 5000 steps of minimization using the steepest descent method keeping only protein backbone atoms fixed to allow protein side chains to relax. After that, the system was equilibrated for 300 ps at constant volume (NVT) and for a further 1 ns using a Langevin piston (NPT)(54) to maintain the pressure while the restraints were gradually released. For every system, three replicates of 1 µs, with different initial velocities, were performed in the NPT ensemble. We applied positional restraints with the force constant of 10 kcal/mol/Å^2^ on the heavy atoms of the M domain during the production runs to avoid drastic rearrangements. The temperature was kept at 310 K and pressure at 1 bar using the Parrinello-Rahman barostat with an integration time step of 2.0 fs. The Particle Mesh Ewald method (PME) (55) was employed to treat long-range electrostatics, and the coordinates of the system were written every 100 ps. Typically, the M1-IgG1 and M1-IgG3 systems are composed of ∼481,152 and ∼1,230,941 atoms in a triclinic water box with a volume of ∼4,925 and ∼12,567 nm3, respectively. The root mean square deviations (RMSD) of the studied complexes from the equilibrated structure were measured on the C$ atoms along simulation time for all the replicates (**Fig. 2**). All systems were fully relaxed after 100 ns. Then, we calculated the residue root mean square fluctuations (RMSF) over the last 900 ns of every simulation (**Fig. 2**) with respect to the average conformation and over the Cα atoms.

#### Hydrogen bonds and salt bridges

The hydrogen bonds (H-bonds) were detected using the HBPLUS algorithm (56). H-bonds are detected between donor (D) and acceptor (A) atoms that satisfy the following geometric criteria: (i) maximum distances of 3.9 Å for D-A and 2.5 Å for H-A, (ii) minimum value of 90° for D-H-A, H-AAA and D-A-AA angles, where AA is the acceptor antecedent. For a given H-bond between residues i and j, interaction strength is computed as the percentage of conformations in which the H-bond is formed between any atoms of the same pair of residues (i and j). Moreover, the salt bridge plugin of the VMD was used to detect all salt bridges at the interface between the Fab domains and the M1 protein (57). The stability of salt bridges was recorded as the percentage of conformations in which the distance between the center of mass of the oxygens in the acidic side chain and the center of mass of the nitrogens in the basic side chain is within the cut-off distance of 3.5 Å.

#### Angles

We measured the angle formed between 1) the Fc domain and the M1 protein, 2) Fc and Fab domains, and 3) Fab1 and Fab2 domains. The vectors used to calculate each angle are defined here:

● *between the Fc domain and the M1 protein*: i) the center of mass of the Fc domain and center of mass of the M1 protein, ii) the center of mass and the N-terminal region of the M1 protein.
● *between the Fc domain and Fab domains*: i) the center of mass of the Fc domain and center of mass of the hinge region, ii) the center of mass of the Fab domain and center of mass of the hinge region
● *between the Fab1 and Fab2 domains*: the center of mass of the hinge region and center of mass of each Fab domain
● *Fab2 domain*: the center of mass of the hinge region and center of mass of the Fab2

The angles were then defined based on the lines that best fit these segments. The results are reported in two scenarios: for all trajectories (**Supp. Fig 3 A-C**) and in dual-Fab binding conformations where the distance between residues at the interface of Fab1 and the M1 protein is less than 30 Å (**Supp. Fig 3 D-F**). The second scenario resulted in 20,074 angles for M1-IgG1, 10,871 angles for M1-IgG3, and 11,306 angles for M1-IgGh simulations.

### Clustering analysis

To characterize representative conformations, we performed a cluster analysis of the MD trajectories using backbone RMSD as the similarity metrics by the GROMOS clustering approach (58) with a cutoff equal to 0.7 nm. The trajectories of replicates were first concatenated, clustering was then performed, and finally, the population of each cluster was calculated. The same analysis was performed only considering the replicates where the dual-Fab binding was observed (replicates 1 and 3 for M1-IgG1, replicate 3 for M1-IgG3, and replicate 1 for IgGh simulations). Clusters with a population of at least 3,000 conformations were colored in shades of green on the plot (**Supp. Fig. 4A**). The results are reported for all MD trajectories of M1-IgG1, M1-IgG3 and IgGh systems (30000, 30000 and 20000 conformations, respectively) and only for dual-Fab binding conformations (20000, 10000 and 10000 conformations, respectively). The three highly populated clusters of IgG1 consist of 9071, 4852, and 3239 conformations. And the top three highly populated clusters of IgG3 consist of 5363, 4195, and 4031 conformations. The representative conformations of the highly populated clusters are shown when dual-Fab binding was observed (**Supp. Fig. 4B-E**).

### Crystallization data analysis

The Ab25 Fab sample was concentrated to 6.75 mg/mL in the buffer of 20 mM HEPES(2-[4-(2-hydroxyethyl)piperazin-1-yl]ethane-1-sulfonic acid), 150 mM NaCl, pH 7.0. A crystallization experiment set up against commercially available screens was performed with a Mosquito crystallization robot (SPT Labtech) using sitting drops with a total volume of 300 nL and different ratios between the protein sample and reservoir solution. Optimization of initial hits gave crystals appearing within a week in 27.50% (w/v) PEG8000, 0.1 M CHES pH 8.5. After growing to full size (50×100×250 µm), the crystals were harvested, cryoprotected in reservoir solution supplemented with 20 % (v/v) glycerol, and flash frozen in liquid nitrogen. Testing for diffraction and data collection was performed at the BioMAX beamline at the MAX IV Laboratory (Lund, Sweden). Diffraction data were collected by fine-slicing with an oscillation range of 0.1°, 360° total. The data were auto-processed with the software EDNA (59) to a resolution of 1.88 Å, and the structure was solved by molecular replacement with PHASER of the PHENIX software suite (60), using PDB entry 6JC2.pdb as a search model for both the heavy and light chain. Electron and difference density maps were manually inspected, and the model was improved using Coot (61) and several rounds of refinement with PHENIX (62). The calculation of Rfree used 5.04 % of the data. Statistics over data and refinement can be seen in **Supplementary Table 1**. The crystallographic data have been deposited to the Protein Data Bank (PDB) with the accession ID: 8C67.

### Statistical analysis

Statistical analysis was performed with Kruskal-Wallis multiple comparisons test with Dunn’s correction test with GraphPad Prism.

## Supplementary Materials

**Movie S1.** Molecular dynamics simulation of IgG1-M1 repeat 1. **Movie S2.** Molecular dynamics simulation of IgG1-M1 repeat 2. **Movie S3.** Molecular dynamics simulation of IgG1-M1 repeat 3. **Movie S4.** Molecular dynamics simulation of IgG3-M1 repeat 1. **Movie S5.** Molecular dynamics simulation of IgG3-M1 repeat 2. **Movie S6.** Molecular dynamics simulation of IgG3-M1 repeat 3. **Movie S7.** Molecular dynamics simulation of IgGh-M1 repeat 1. **Movie S8.** Molecular dynamics simulation of IgGh-M1 repeat 2. **Movie S9.** Molecular dynamics simulation of IgGh-M1 repeat 3.

## Acknowledgments

This work was funded by the Swedish Research Council (Vetenskapsrådet, PN, JM), the Knut and Alice Wallenberg Foundation (PN, OS, LM, JM), the Institut Pasteur (Pasteur-Cantarini-Roux fellowship to Y.K.). This work was granted access to the HPC resources of IDRIS under the allocation 2021-A0110711500 made by GENCI (granted to Y.K. and M.N.). HK was supported by the French Agence Nationale de la Recherche (ANR), under grant ANR-22-CPJ2-0075-01.

## Author Contributions

A.I., YK, HK, JM, WB, OS, LM and PN. designed research; A.I., YK, EB, SW, HK, MNy, BO, LH and DT, performed research; A.I, YK, EB, HK, MNy and LH. analyzed data; A.I., YK, and P.N. wrote the paper. All authors contributed to reading and editing the final manuscript.

---

**Supplementary Figure 1.**
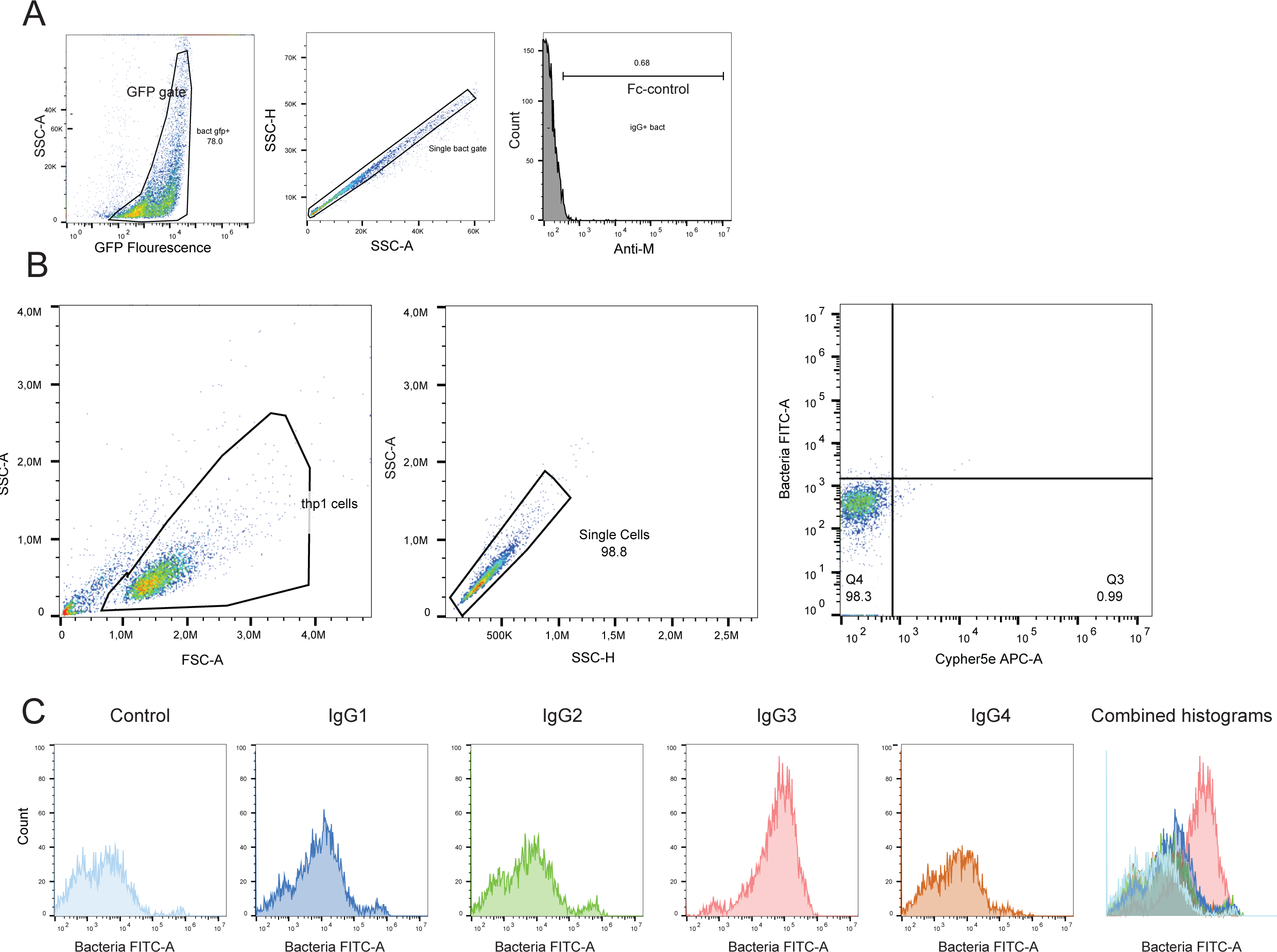
Gating strategies for flow cytometry-based binding and phagocytosis assays. **A** Gating strategy for bacteria. Bacteria were gated for granularity (SSC-A) and GFP fluorescence (FITC-A). Thereafter single cell gating was done to exclude duplicate events based on granularity height (SSC-H) and area (SSC-A). To assess IgG binding to bacteria, we used a low concentration of Fc-control antibody (0.5 ug/mL) to set the gate for secondary binding (APC-A). **B** THP-1 cells were gated for by size and granularity. Thereafter single cell gating was done to eliminate duplicate events. To determine the level of internalization and association we used a negative control with cells only. Cells internalized with bacteria would be both positive for FITC (Oregon green stained bacteria) but also APC (CypHer5e pH-sensitive dye). Cells are associated with bacteria when only positive in the FITC-channel. **C** the amount of bacteria (FITC) being phagocytosed by the population of THP-1 cells in the single cell gate. From left to right histograms of control, IgG1, IgG2, IgG3, IgG4 is shown ending with a combined histogram of all five plots.

**Supplementary Figure 2.**
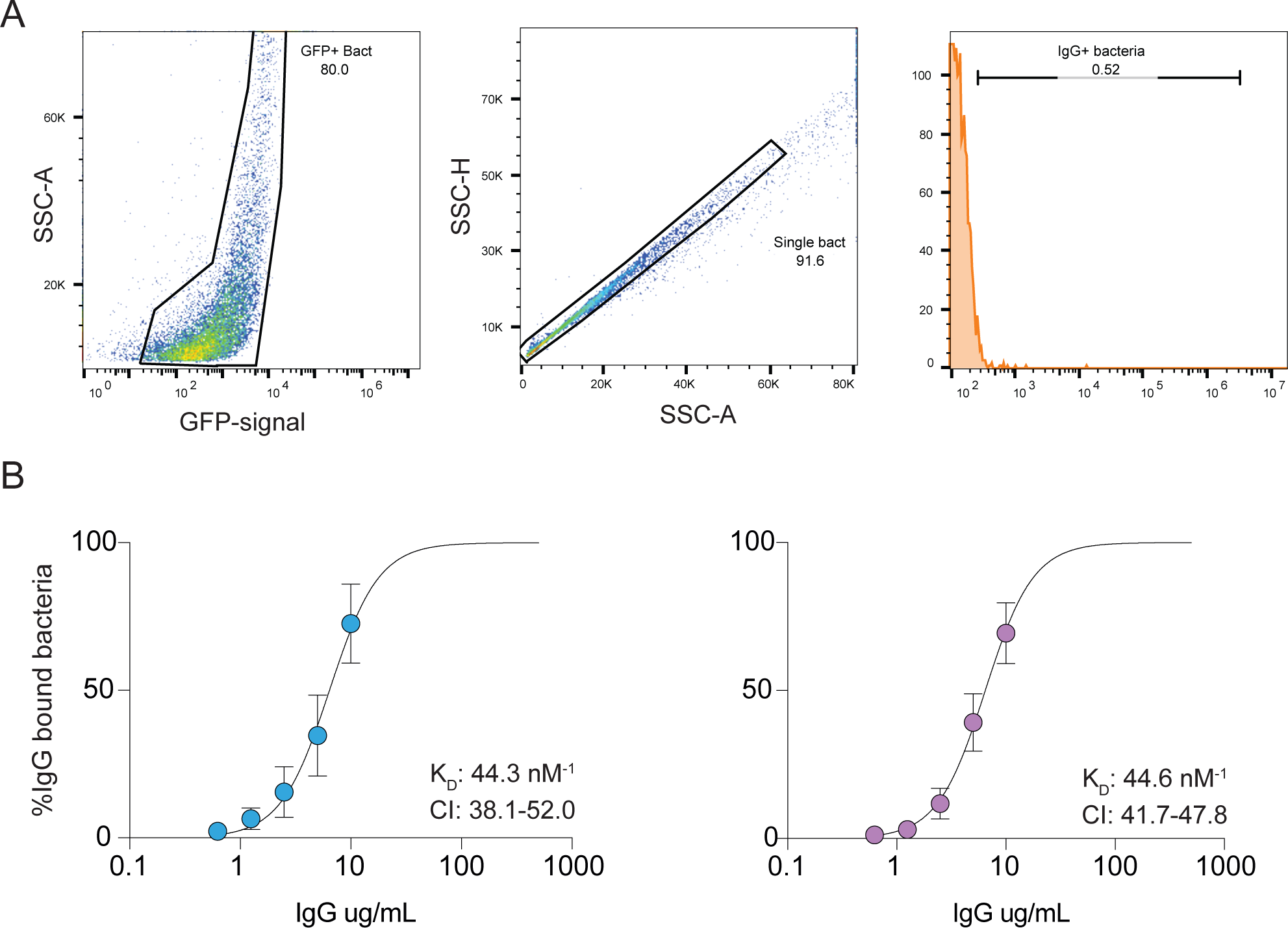
Gating strategies for flow cytometry-based binding and phagocytosis assays. **A** Gating strategy for bacteria. Bacteria were gated for granularity (SSC-A) and GFP fluorescence (FITC-A). Thereafter single cell gate was done to exclude duplicate events based on granularity height (SSC-H) and area (SSC-A). To assess IgG binding to bacteria directly conjugated IgG with Alexa 647 (APC-A) and used unstained bacteria to set the gate. **B** Affinity curves for the directly conjugated antibodies. The data points are the mean value of four independent experiments with biological replicates. The KD values are shown with the 95% confidence interval indicated.

**Supplementary Figure 3.**
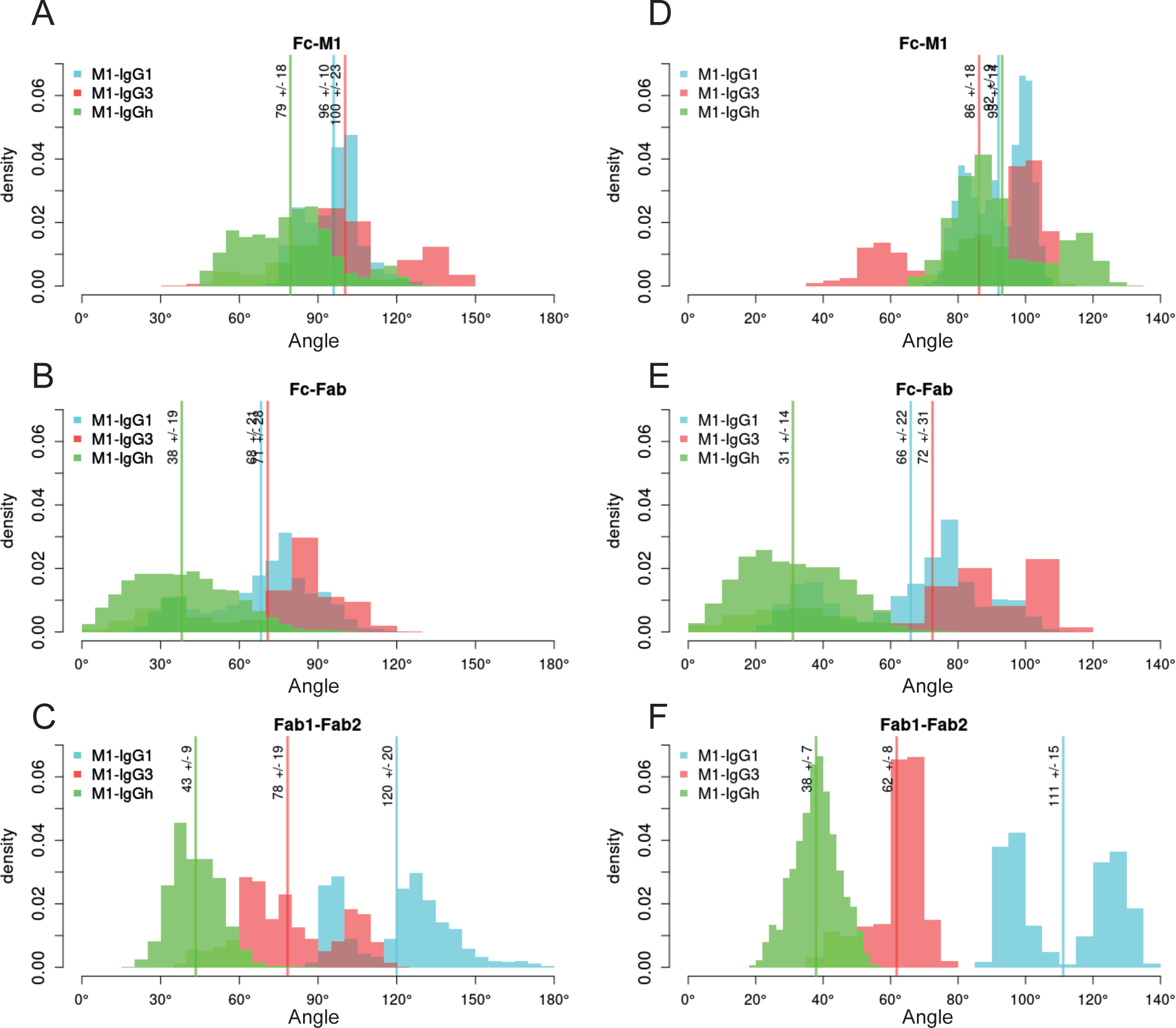
Changes of angles along the MD simulations for M1-IgG1, M1-IgG3 and M1-IgGh systems. The Y-axis shows the probability densities of the MD conformations for that specific angle while the X-axis shows the angle in degrees. A-C are the values reported over all the MD trajectories, while D-F are when the Fab domains are within 30 Å of the M1 protein. A and D show the angles between the Fc domain and M1. B and E show the angles formed between the Fc and Fab domains. C and F show the angles formed between Fab 1 and Fab 2. The IgG1-M1 system is highlighted in blue, red for IgG3-M1 and green for IgGh-M1 respectively.

**Supplementary Figure 4.**
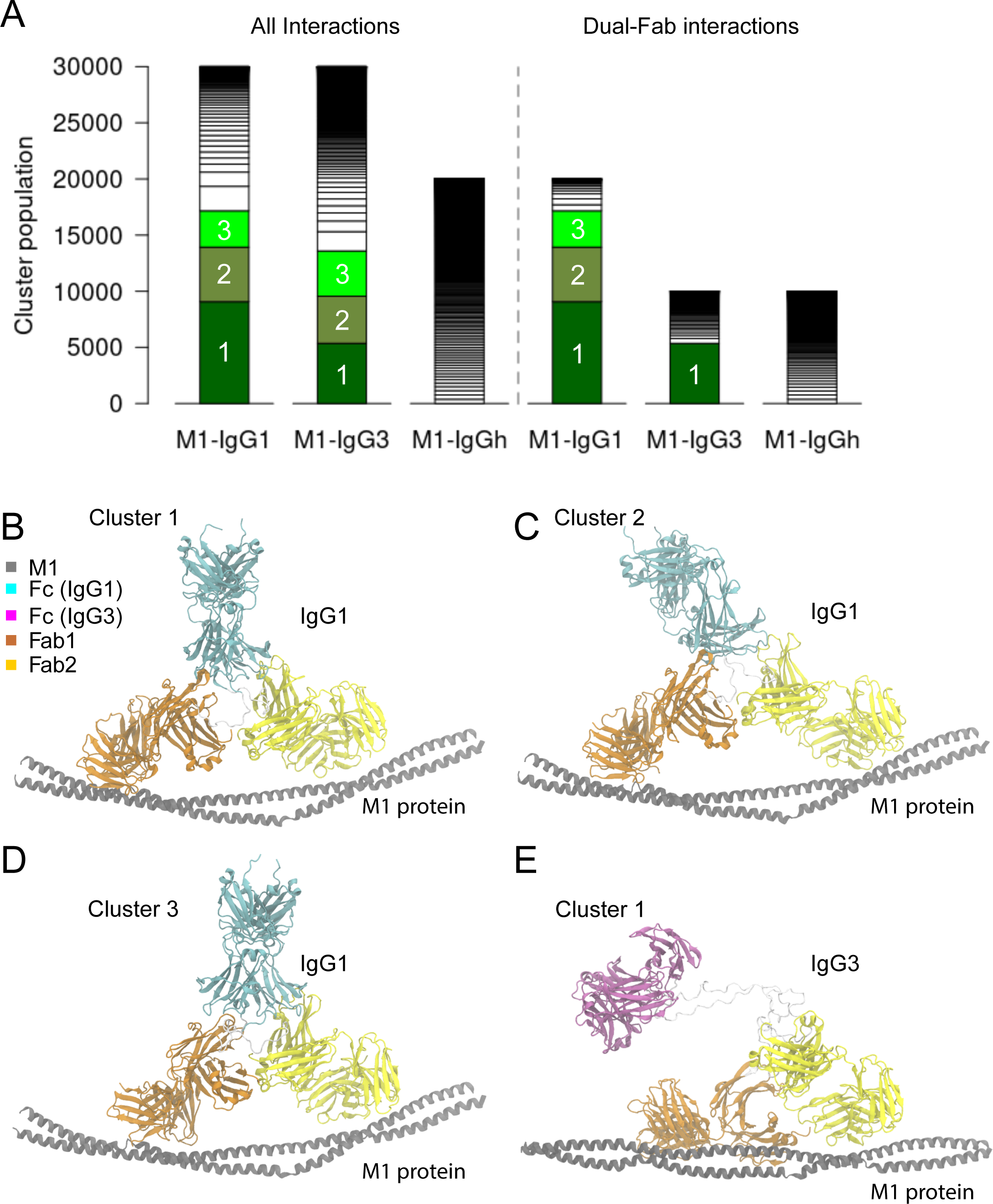
Clustering of MD confirmations for IgG1-M1, IgG3-M1 and IgGh-M1 systems. A The results are reported over all MD trajectories and only dual-Fab binding conformations. Highly populated clusters (with at least 3000 conformations) are colored in shades of green.The representative dual-Fab binding conformation for cluster 1 in B, cluster 2 in C and cluster 3 in D for M1-IgG1 and cluster 1 for M1-IgG3 in E.

**Supplementary Figure 5.**
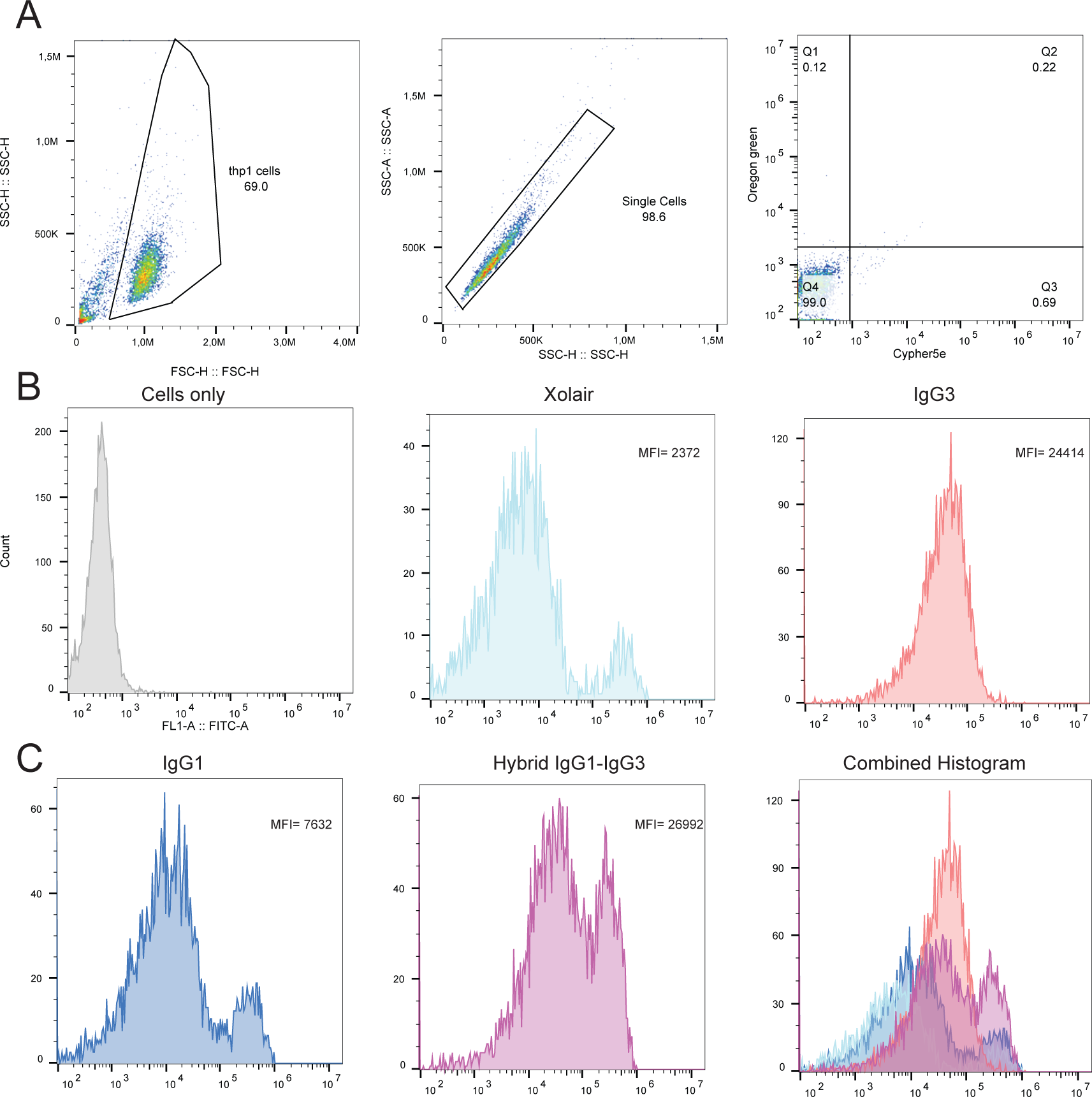
Gating strategies for flow cytometry phagocytosis assays. THP-1 cells were gated for size and granularity. Thereafter a single cell gate was done to eliminate inclusion of duplicate events. To determine the level of internalization and association we used a negative control with cells only. Cells with internalized bacteria would be both positive for FITC (Oregon green stained bacteria) but also APC (Cypher5e pH-sensitive dye stained). Cells are associated with bacteria when only positive in the FITC-channel. C Highlights the amount of bacteria (FITC) being phagocytosed by the population of THP-1 cells in the single cell gate. From left to right the histograms of cells only, Xolair, IgG3, IgG1, Hybrid IgG and a combined histogram of all five. The Y-axis shows count and the x-axis MFI (FITC). The median FITC-A signal is highlighted in the graph in the top left corner.

**Supplementary Table 1.** Crystallography data table over data used in structure determination. Data in parenthesis correspond to the highest resolution shell.

| Data collection | Ab25 |
| --- | --- |
| Space group | P 1 21 1 |
| Cell dimensions<br>a, b, c (Å) | 86.90 71.89 146.01 |
| $\alpha$, $\beta$, $\gamma$ (°) | 90.00 90.10 90.00 |
| Wavelength (Å) | 0.97625 |
| Resolution (Å) | 48.67-1.88 (1.95-1.88) |
| Rmerge | 0.119 (1.107) |
| Number of observations | 1015596 (96059) |
| Number of unique observations | 146332 (14268) |
| Mean $I/\sigma(I)$ | 11.7 (1.9) |
| Completeness (%) | 99.9 (99.7) |
| Multiplicity | 6.9 (6.7) |
| CC(1/2) | 0.998 (0.835) |
| **Refinement** |  |
| Resolution (Å) | 44.152-1.880 |
| $R_{\text{work}}/R_{\text{free}}$ | 0.2050/ 0.2546 |
| Number of reflections | 146123 |
| $R_{\text{free}}$ test set, reflections (%) | 5.04 |
| No. of non-H atoms | 14184 |
| *R.m.s. deviation from ideal geometry* |  |
| bonds (Å) | 0.008 |
| angles (°) | 1.090 |
| Average B, all atoms (Å<sup>2</sup>) | 39.4 (range 14.0–115.96) |
| *Ramachandran plot* |  |
| Favoured regions (%) | 96.89 |
| Allowed regions (%) | 2.76 |
| Outliers (%) | 0.35 |
| MolProbity clash score | 4.14 |

**Supplementary Table 2.**
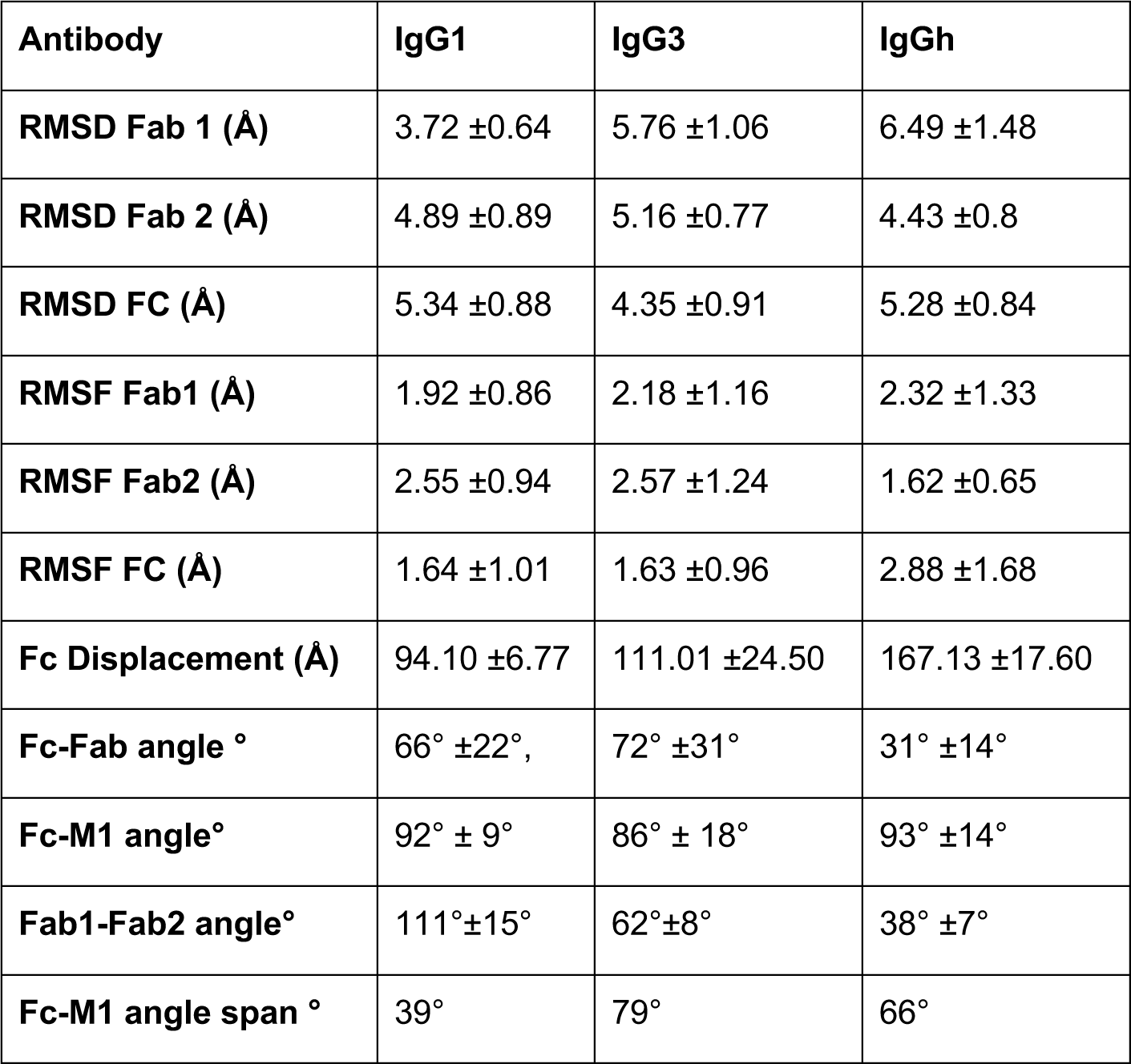
Data summary from MD simulations.

| Antibody | IgG1 | IgG3 | IgGh |
| --- | --- | --- | --- |
| RMSD Fab 1 (Å) | 3.72 ±0.64 | 5.76 ±1.06 | 6.49 ±1.48 |
| RMSD Fab 2 (Å) | 4.89 ±0.89 | 5.16 ±0.77 | 4.43 ±0.8 |
| RMSD FC (Å) | 5.34 ±0.88 | 4.35 ±0.91 | 5.28 ±0.84 |
| RMSF Fab1 (Å) | 1.92 ±0.86 | 2.18 ±1.16 | 2.32 ±1.33 |
| RMSF Fab2 (Å) | 2.55 ±0.94 | 2.57 ±1.24 | 1.62 ±0.65 |
| RMSF FC (Å) | 1.64 ±1.01 | 1.63 ±0.96 | 2.88 ±1.68 |
| Fc Displacement (Å) | 94.10 ±6.77 | 111.01 ±24.50 | 167.13 ±17.60 |
| Fc-Fab angle ° | 66° ±22°, | 72° ±31° | 31° ±14° |
| Fc-M1 angle° | 92° ± 9° | 86° ± 18° | 93° ±14° |
| Fab1-Fab2 angle° | 111°±15° | 62°±8° | 38° ±7° |
| Fc-M1 angle span ° | 39° | 79° | 66° |

